# Specific presynaptic functions require distinct Drosophila Ca_v_2 splice isoforms

**DOI:** 10.1101/2024.06.14.598978

**Authors:** Christopher Bell, Lukas Kilo, Daniel Gottschalk, Jashar Arian, Lea Deneke, Hanna Kern, Christof Rickert, Oliver Kobler, Julia Strauß, Martin Heine, Carsten Duch, Stefanie Ryglewski

**Author notes:** these authors contributed equally (Christopher Bell, Lukas Kilo).

## Abstract

The multiplicity of neural circuits that accommodate the sheer infinite number of computations conducted by brains requires diverse synapse and neuron types. At many vertebrate synapses release probability and other aspects of presynaptic function are tuned by different combinations of Ca_v_2.1, Ca_v_2.2, and Ca_v_2.3 channels. By contrast, most invertebrate genomes contain only one *Ca*_v_*2* gene. The one Drosophila Ca_v_2 homolog, cacophony (cac), localizes to presynaptic active zones (AZs) to induce synaptic vesicle release. We hypothesize that Drosophila cac functional diversity is enhanced by two specific exon pairs that are mutually exclusively spliced and not conserved in vertebrates, one in the voltage sensor and one in the intracellular loop containing the binding site(s) for Ca_β_ and G-protein βγ subunits. We test our hypothesis by combining opto– and electrophysiological with neuroanatomical approaches at a fast glutamatergic model synapse, the Drosophila larval neuromuscular junction. We find that alternative splicing in the voltage sensor affects channel activation voltage and is imperative for normal synapse function. Only the isoform with the higher activation voltage localizes to the presynaptic AZ and mediates evoked release. Removal of these cac splice isoforms renders fast glutamatergic synapses non-functional. By contrast, alternative splicing at the other alternative exon that encodes the intracellular loop between the first and the second homologous repeats does not affect cac presynaptic AZ localization, but it tunes multiple aspects of presynaptic function. While expression of one exon yields normal transmission, expression of the other exon reduces channel number in the AZ and thus release probability. This also abolishes presynaptic homeostatic plasticity. Moreover, reduced channel number upon selective exon excision increases paired pulse ratios and the variability of synaptic depression during low frequency stimulation trains (1 and 10 Hz), and thus affects short term plasticity. Effects on short term plasticity can be rescued by increasing the external calcium concentration to match release probability to control. In sum, in Drosophila alternative splicing provides a mechanism to regulate different aspects of presynaptic functions with only one *Ca*_v_*2* gene.

## Introduction

Information transfer at fast chemical synapses requires action potential triggered calcium influx through voltage gated calcium channels (VGCCs) into the presynaptic terminal, which in turn initiates synaptic vesicle (SV) release through a complex cascade of biophysical and biochemical reactions (Südhof 2013; 2014; Dittman, Ryan 2019). The millisecond temporal precision of SV release upon the arrival of an action potential in the axon terminal requires clustering of the VGCCs in a spatially restricted presynaptic specialization, named the presynaptic active zone (AZ; Eggermann et al. 2011; Südhof 2012; Van Vactor, Sigrist 2017; Dolphin, Lee 2020; Emperador-Melero, Kaeser 2020). At the AZ, protein-protein interactions ensure a nanoscale coupling of VGCCs to SVs in the readily releasable pool (Kittel et al. 2006), an arrangement that supports efficient and fast neurotransmission (Eggermann et al. 2011).

Despite these common organizational principles, fast chemical synapses are highly heterogeneous to accommodate the diverse computational requirements of different types of neurons and brain circuits (Moser et al. 2023; Zhang et al. 2022). One means of synapse diversification is to organize VGCCs heterogeneously at the AZ to modify either their nanoscale spatial relation to the SVs ready for release (Dittman, Ryan, 2019; Rebola et al. 2019), or the profiles of the calcium dynamics that shape release probability (P_r_; Zhang et al. 2022). The calcium dynamics in nano-or microdomains are affected by VGCC number, clustering, and properties, and are thus strongly dependent on calcium channel subtype.

The vertebrate genome contains 10 genes encoding the α_1_-subunit of VGCCs that fall into three families (Ca_v_1, Ca_v_2, Ca_v_3; Dolphin, 2009). With the exception of Ca_v_1 triggering release at retinal ribbon and auditory brain stem synapses (Moser et al. 2020), Ca_v_2 usually mediates evoked release at other mammalian synapses. The three subtypes of Ca_v_2 exhibit different biophysical properties (Sheng et al. 2012), which may explain their different contribution to SV release at a given synapse (Li et al. 2007). Ca_v_2.1 and Ca_v_2.2 trigger SV release at most synapses and Ca_v_2.3 likely make only a small contribution (Dietrich et al. 2003). However, all three Ca_v_2 subtypes may co-localize to the same synapse and contribute to SV release (Takahashi, Momiyama 1993; Wheeler et al. 1994; Wu et al. 1998), and different types of mammalian synapses can utilize either only one subtype or combinations of Ca_v_2 subtypes (reviewed in Zhang et al. 2022).

The Drosophila genome contains only 3 genes encoding α_1_-subunits of VGCCs, each one homologous to one vertebrate Ca_v_ family (Littleton, Genetzky 2000). Therefore, the joint functions of mammalian Ca_v_2.1, Ca_v_2.2, and Ca_v_2.3 are covered by only one Drosophila gene, namely *Dmca1A* (also named *cacophony* or *nightblindA*, Smith et al. 1996; 1998). Cacophony (cac) is, in fact, essential for fast synaptic transmission in Drosophila (Kawasaki et al. 2002) and loss of cac cannot be compensated for by the Drosophila counterparts of Ca_v_1 or Ca_v_3 (Krick et al. 2021). This suggests that Drosophila lacks the combinatorial logic of employing different blends of Ca_v_2.1, Ca_v_2.2, and Ca_v_2.3 for fine-tuning of P_r_ as present in mammals. However, the Drosophila *cac* locus contains two mutually exclusive alternative splice sites that are not present in the vertebrate *Ca*_v_*2* gene family, one in the voltage sensor in the fourth transmembrane domain of the first homologous repeat (IS4) and one in the intracellular linker between the first and the second homologous repeats (I-II). Alternative splicing at these mutually exclusive sites may provide mechanisms to fine tune action potential triggered calcium dynamics at presynaptic AZs by adjusting the numbers, nanoscale localization and properties of presynaptic cacophony VGCCs. We test this hypothesis by employing the CRISPR-Cas9 technology to remove mutually exclusive exons at either the *IS4* or the *I-II* site and test the resulting functional consequences at the Drosophila neuromuscular junction (NMJ), a well-established model for presynaptic function of a fast glutamatergic synapse (Atwood, Karunanithi 2002; Harris, Littleton 2015).

We find that alternative splicing at the *IS4* site affects cac voltage activation and only one of both mutually exclusive splice events allows for presynaptic AZ localization and thus evoked synaptic transmission at a fast glutamatergic synapse. By contrast, alternative splicing at the *I-II* site does not affect presynaptic AZ localization, but it affects P_r_, short term plasticity, and presynaptic homeostatic plasticity.

## Results

*Cacophony* (*cac*) is located on the X-chromosome and contains multiple alternative splice sites but only two that are mutually exclusive (Fig. 1A, see also enlarged schematic of the cac regions housing these two mutually exclusive exon pairs (green boxes); cac schematic in Fig. 1B). The first one is located in the fourth transmembrane domain of the first homologous repeat (IS4), and thus affects the voltage sensor. Both alternative *IS4* exons are 99 bp long and encode 33 amino acids (AAs). The polypeptide encoded by *IS4A* (first alternative exon of the *IS4* locus) differs from that of *IS4B* in 13 AAs and contains one additional positively charged AA. The second mutually exclusive exon pair encodes part of the intracellular linker between homologous repeats I and II (*I-II*; Fig. 1B), and is thus predicted to affect binding sites for calcium channel β subunits (Ca_β_) and G-protein βγ subunits (G_βγ_). Both alternative *I-II* exons are 117 bp long and encode 39 AAs. The polypeptide encoded by *I-IIA* (first alternative exon of the *I-II* locus) differs from that of *I-IIB* in 23 AAs. Sequence analysis strongly suggests that I-IIB contains both, a Ca_β_ as well as a G_βγ_ binding site (AID: α-interacting domain) because the binding motif QXXER is present. In mouse Ca_v_2.1 and Ca_v_2.2 channels the sequence is QQIER, while in Drosophila cacophony I-IIB it is QQLER. In the alternative I-II linker I-IIA, this motif is not present, strongly suggesting that G_βγ_ subunits cannot interact at the AID. However, as already suggested by Smith et al. (1998), also based on sequence analysis, Ca_β_ should still be able to bind, although possibly with a lower affinity. We employ CRISPR-Cas9 to excise one of both alternative exons either at the *IS4* or the *I-II* loci (Fig. 1C). Genomic removal of *IS4A* results in fly strains that contain only the *IS4B* exon and are named *ΔIS4A* (Fig. 1C). Accordingly, genomic removal of the *I-IIA* exon results in flies with *I-IIB* only that are named *ΔI-IIA* (Fig. 1C). To analyze the functions of mutually exclusive exon variants, we create exon-out fly strains with removal of one exon at a time of each of the four exons (*IS4A*, *IS4B*, *I-IIA*, *I-IIB*). Excision of the respective exon is always confirmed by sequencing (see methods). Fly strains with first chromosomes that carry exon excisions are cantonized by 10 generations of backcrossing into Canton S wildtype flies. Exon-out strains are produced from either wildtype (Canton S) or white mutant flies (*w^1118^*, see methods) as well as from fly strains with previously introduced N-terminal fluorophore tagging of the *cac* gene (Gratz et al. 2019). Neither on-locus super folder GFP (sfGFP) tagging, nor on-locus tagging of the endogenous cacophony channel with mEOS4b (Ghelani et al. 2023) impairs cac localization at presynaptic AZs or synaptic function at the larval Drosophila NMJ. In this study, sfGFP-tagged (Fig. 1B) exon-variants serve analysis of channel localization and mEOS4b tagged (Fig. 1B) ones serve to count channel number in AZs, and untagged fly strains serve to control for possible effects of the fluorescent tags.

**Figure 1.**
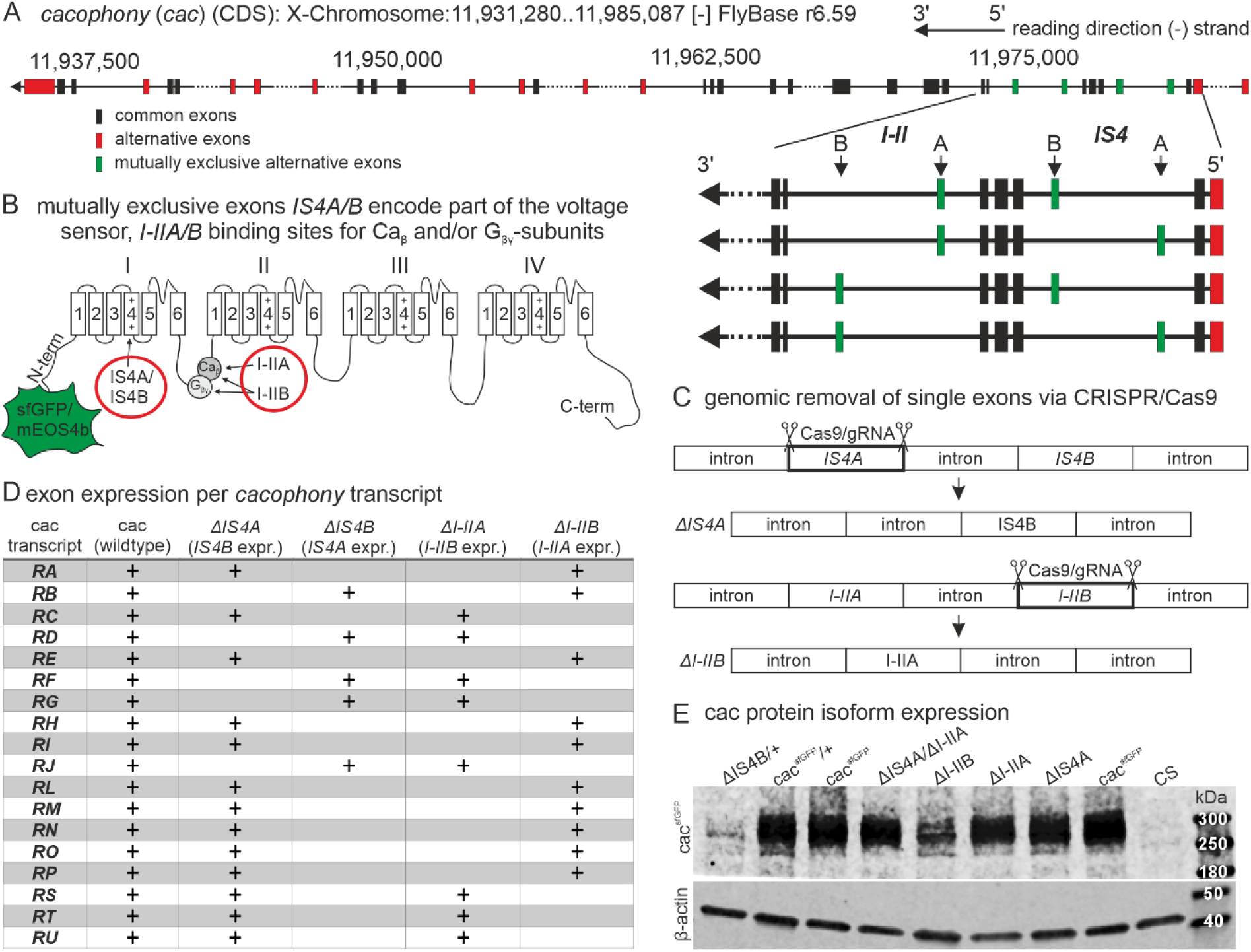
Cacophony alternative splicing gives rise to different protein isoforms. **(A)** *Cacophony* is located on the non-coding strand of the X-chromosome between positions 11,931,280 and 11,985,087 (Flybase version r6.59). Reading direction is indicated on the top right (arrow). Horizontal black lines indicate introns, larger introns are broken by dots. Black boxes indicate exons shared by all *cacophony* splice variants, red boxes indicate alternative exons, green boxes represent exons that are mutually exclusively spliced. Enlarged area on the right emphasizes mutually exclusive exon pairs *IS4A/B* and *I-IIA/B*. **(B)** Schematic of cacophony with N-terminal sfGFP or mEOS4b tag and two exon pairs that are spliced mutually exclusively. *IS4A* and *IS4B* exons encode isoforms of the 4^th^ transmembrane domain (S4) and thus part of the voltage sensor of the first homologous repeat (I) of the calcium channel, while *I-IIA* and *I-IIB* give rise to two versions of the intracellular linker between homologous repeats I and II. I-IIA contains a non-well conserved binding site for voltage gated calcium channel β-subunits (Ca_β_) whereas I-IIB gives rise to a conserved Ca_β_-binding site as well as a binding site for G-protein βγ-subunits (G_βγ_). **(B)** Genomic removal of one of the mutually exclusively spliced exons of one or two exon pairs by CRISPR/Cas9 mediated double strand breaks in the germ line results in exon out mutants by imprecise excision of the cut exons and cell-intrinsic DNA repair. **(C)** *Cacophony* gives rise to 18 annotated transcripts (RA-RU, left). Multiple variants express the same mutually exclusive exons but differ with respect to expression of other alternatively spliced but not mutually exclusive exons. Removal of the mutually exclusive exons *IS4A/IS4B* and/or *I-IIA/I-IIB* allows expression of fewer *cacophony* splice variants. Transcript variants that are possible upon exon excision are marked by +. **(D)** Western Blots reveal expression of GFP-tagged cacophony protein (cac^sfGFP^) for all excision variants at the expected band size of ∼240 kDa (cacophony) plus ∼30 kDa (sfGFP, top), while no band can be detected in the Canton S wildtype (CS, top, right) that does not express cac^sfGFP^. 10 adult brains were used of hemizygous males for all genotypes except for *ΔIS4B* which is homo-/hemizygous lethal. For *ΔIS4B* 20 brains of heterozygous females and a heterozygous cac^sfGFP^ control were used (F1 females from cross with Canton S wildtype flies(+)). β-actin was used as loading control (bottom). *ΔIS4B* shows weak expression (top, left), while all other exon out variants express strongly, although *ΔI-IIB* shows somewhat weaker expression (top, middle).

Of the 18 annotated *cacophony* transcripts in Drosophila (Flybase, r6.59), 13 remain upon removal of *IS4A*, 5 upon removal of *IS4B*, 8 upon removal of *I-IIA*, and 10 upon removal of *I-IIB* (Fig. 1D). For each mutually exclusive exon, we have created multiple exon-out fly strains. Each exon removal induces reproducible phenotypes (see methods). Removal of *IS4B* is embryonic lethal as confirmed in all CRISPR-Cas9 mediated excisions (n=12), both before and after crossing out into a wildtype background. Removal of each of the three other mutually exclusive exons results in viable fly strains, Drosophila larvae without any obvious deficits, before and after crossing out into a wildtype background, but with distinct behavioral phenotypes in adult flies (the latter are not further addressed in this study). Western blots with GFP tagged cac channels test whether the channel protein is expressed in the CNS of all exon-out fly strains (Fig. 1E). *ΔIS4B* flies are homozygous lethal and are thus used heterozygously over untagged wildtype channels that contain all exons for Western blot analysis. Brain homogenate from sfGFP tagged controls and from all exon-out fly strains with sfGFP tagged cac channels yield a band at roughly the expected size of 250-270 kDa (Drosophila cac isoforms range from 212 to 242 kDa and sfGFP is 27 kDa), whereas no band is detected in control brains without tagged cac (Fig. 1E, right lane). Protein level in *ΔIS4B* flies (first band from left, Fig. 1E) is lower as compared to heterozygous controls with all isoforms (second band from left, Fig. 1E). Similarly, protein level in *ΔI-IIB* flies (fifth band from left, Fig. 1E) is lower as compared to homozygous controls with all isoforms (third and eighth band from left, Fig. 1E), thus indicating that cac isoforms containing IS4B and I-IIB are normally abundantly expressed. By contrast, homozygous removal of *IS4A* or of *I-IIA* does not cause obvious differences in expression levels as compared to homozygous sfGFP tagged controls (Fig. 1E). Taken together, CRISPR-Cas9 is successfully employed to produce all possible distinct exon excisions at the mutually exclusive splice sites of Drosophila *cacophony*. This now allows for analysis of *cac* exon specific functions.

We next analyze the functional consequences of the removal of each of the mutually exclusive exons at the *IS4* and at the *I-II* loci for cac channel localization, channel number, and channel function at motoneuron presynaptic terminals of the larval Drosophila neuromuscular junction (NMJ).

### IS4B localizes to presynaptic active zones and is required for evoked synaptic transmission

Drosophila cac channels interact with the scaffold protein bruchpilot (brp) to localize to presynaptic AZs (Kittel et al. 2006; Ghelani et al. 2023). Immunohistochemical triple label at the larval Drosophila NMJ tests whether either one of the mutually exclusive exons at the *IS4* locus is required for correct presynaptic cac localization. On the level of CLSM, we define correct cac localization in presynaptic AZs at the larval NMJ as cac puncta that overlap with brp puncta but no cac label anywhere else in the presynaptic bouton. Motoneuron axon terminals on larval muscles 6 and 7 (M6/7) are labeled with HRP (Figs. 2A-C, right column, blue), cac^sfGFP^ by α-GFP immunolabel (Figs. 2A-C, left column, green), and AZs by α-brp immunohistochemistry (Figs. 2A-C, second column, magenta). For a better visualization of label in AZs, figures 2Ai-Ci show selective enlargements of single boutons with all labels. In controls, cac^sfGFP^ channels with full isoform diversity strictly co-localize with brp in presynaptic AZs (Fig. 2A, cac and brp overlay, 3^rd^ column) as previously reported (Gratz et al. 2019). The same is the case for cac channels that contain IS4B but lack IS4A *(ΔIS4A*, Fig. 2B). By contrast, cac channels that contain IS4A but lack IS4B (*ΔIS4B*) do not localize to presynaptic AZs (Fig. 2C). On the level of CLS microscopy we define strict AZ localization as cac puncta clearly overlapping with brp puncta but no cac label anywhere else in the synaptic bouton. Importantly, upon excision of *IS4B*, cac channels are not only absent from AZs, but immunolabel for tagged IS4A channels cannot be detected above background anywhere in the axon terminals (Fig. 2Ci, see also below Fig. 3A, bottom). Since *ΔIS4B* is homozygous lethal, heterozygous animals are used for triple labels of tagged IS4A channels, the AZ marker brp, and motoneuron terminals on M6/7. Axon terminal shape and brp label in AZs are qualitatively normal, but the label for cac^IS4A^ (=*ΔIS4B*) channels is absent (Fig. 2Ci). Accordingly, the Pearson’s correlation coefficient of IS4A and brp is ∼0.2 (Fig. 2D), which is considered negligible (Mukaka 2012), whereas cac and IS4B (=*ΔIS4A*) channels show a significant Pearson’s correlation coefficient as previously reported for controls (Krick et al. 2021). In addition, quantification of brp labeling intensity did not show any difference between genotypes, not even in the absence of cac upon excision of *IS4B* (data not shown). These data indicate that the *IS4B* exon is required for targeting/localizing cac channels within the AZ.

**Figure 2.**
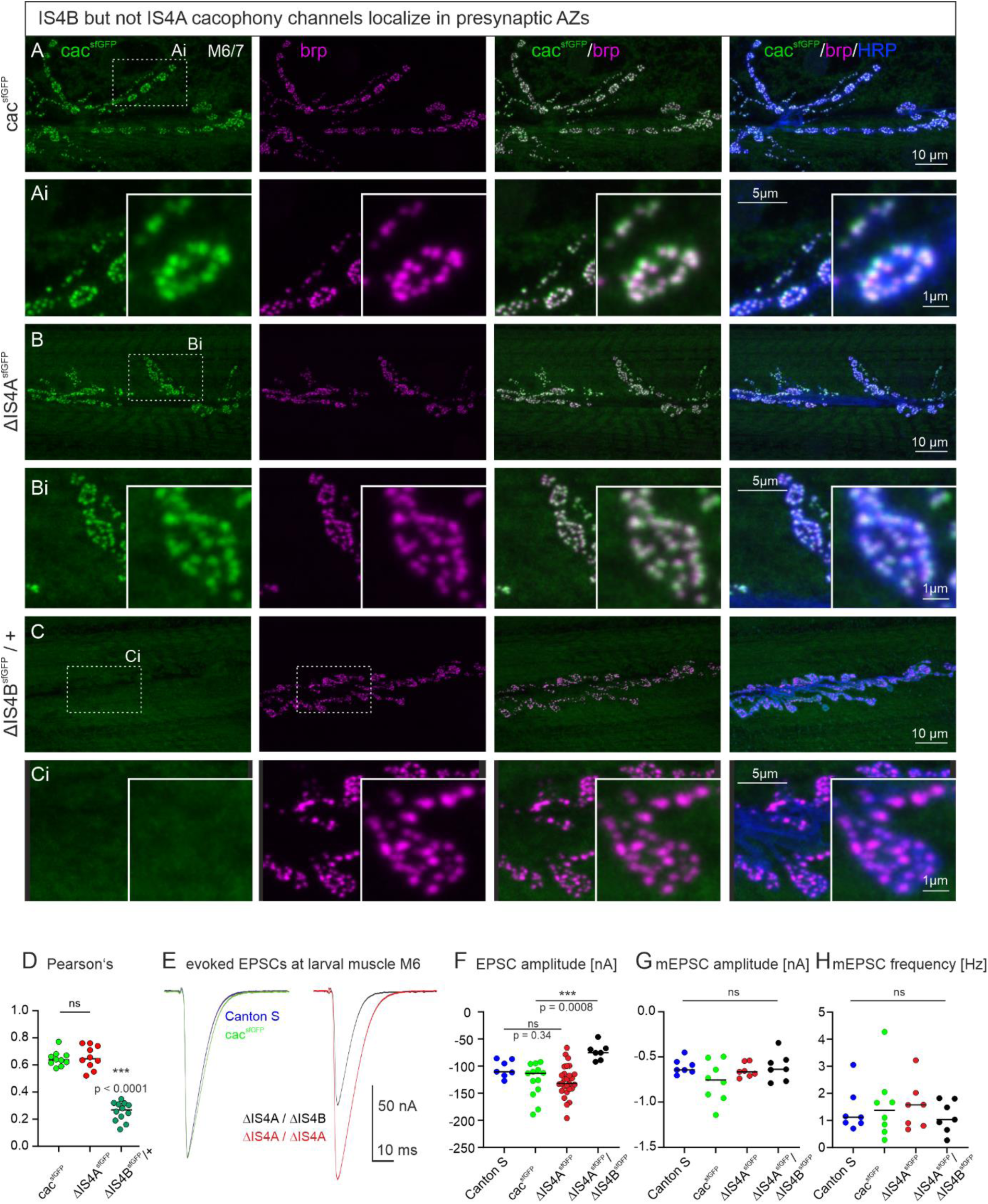
The IS4B exon is required for cacophony localization to AZs and for evoked synaptic transmission. **(A-C)** Representative confocal projection views of triple labels for GFP tagged cac channels (green), the AZ marker brp (magenta), and HRP to label axonal membrane (blue) as well as enlargements of the area marked by dotted rectangles (Ai-Ci). Label was done in control animals with all cac exons (cac^sfGFP^, top row, **A, Ai**), in animals with selective excision of either the alternative exon *IS4A (ΔIS4A^sfGFP^*, middle rows, **B, Bi**), or the alternative exon *IS4B* (*ΔIS4B^sfGFP^*, bottom row, **C, Ci**). Excision of *IS4B* is embryonic lethal, so that localization analysis was conducted in heterozygous animals (*ΔIS4B^sfGFP^/+*). The gross morphology of the neuromuscular junctions (muscle fibers, bouton numbers and sizes, AZ numbers) was similar in all three genotypes. GFP tagged cac channels localize to AZs **(A, Ai)** as previously reported (Gratz et al. 2019; Krick et al. 2021). Excision of the *IS4A* exon does neither impact cac AZ localization nor labeling intensity **(B, Bi)**. By contrast, upon excision of the *IS4B* exon, no cac label is detected **(C, Ci)**. **(D)** Pearson’s correlation analysis of cac and brp in cac^sfGFP^ (green), *ΔIS4A* (red), and heterozygous *ΔIS4B* (dark green) animals reveals correlation coefficients of ∼0.6 for cac^sfGFP^ and *ΔIS4A* but not for *ΔIS4B* (Pearson’s correlation coefficient ∼0.26, which is significantly different from cac^sfGFP^ control (p<0.001, ANOVA with Dunnett’s post hoc test, comparison only against control) and considered negligible. This is in line with absent label in *ΔIS4B* animals (C, Ci). **(E, F)** Representative traces of evoked synaptic transmission as recorded in TEVC from muscle fiber 6 upon extracellular stimulation of the motor nerve from a wildtype control animal (CS, blue), an animal cac^sfGFP^ (green), and animals with cac^sfGFP^ and either *IS4A* exon excision (*ΔISA4*, red) or transheterozygous female animals with *IS4B* excision over *IS4A* excision (*ΔISAB/ΔISA4*, black). **(E)** Excitatory postsynaptic currents (EPSCs) are similarly shaped between CS control (blue) and animals expressing cac^sfGFP^ (green), and **(F)** EPSC amplitudes are not statistically different (p=0.34, two sided Tukey’s multiple comparison test). In animals with homozygous *IS4A* exon excision (red), EPSC amplitude is slightly but not significantly increased (p=0.18, two sided Tukey’s multiple comparison test). In transheterozygous animals with *IS4A* excision on one chromosome and *IS4B* excision on the other one, EPSC amplitude is significantly decreased (p=0.0008), two sided Tukey’s multiple comparison test). **(G)** Quantal size (mEPSC amplitude) and spontaneous release frequency **(H)** show no significant difference between genotypes.

**Figure 3.**
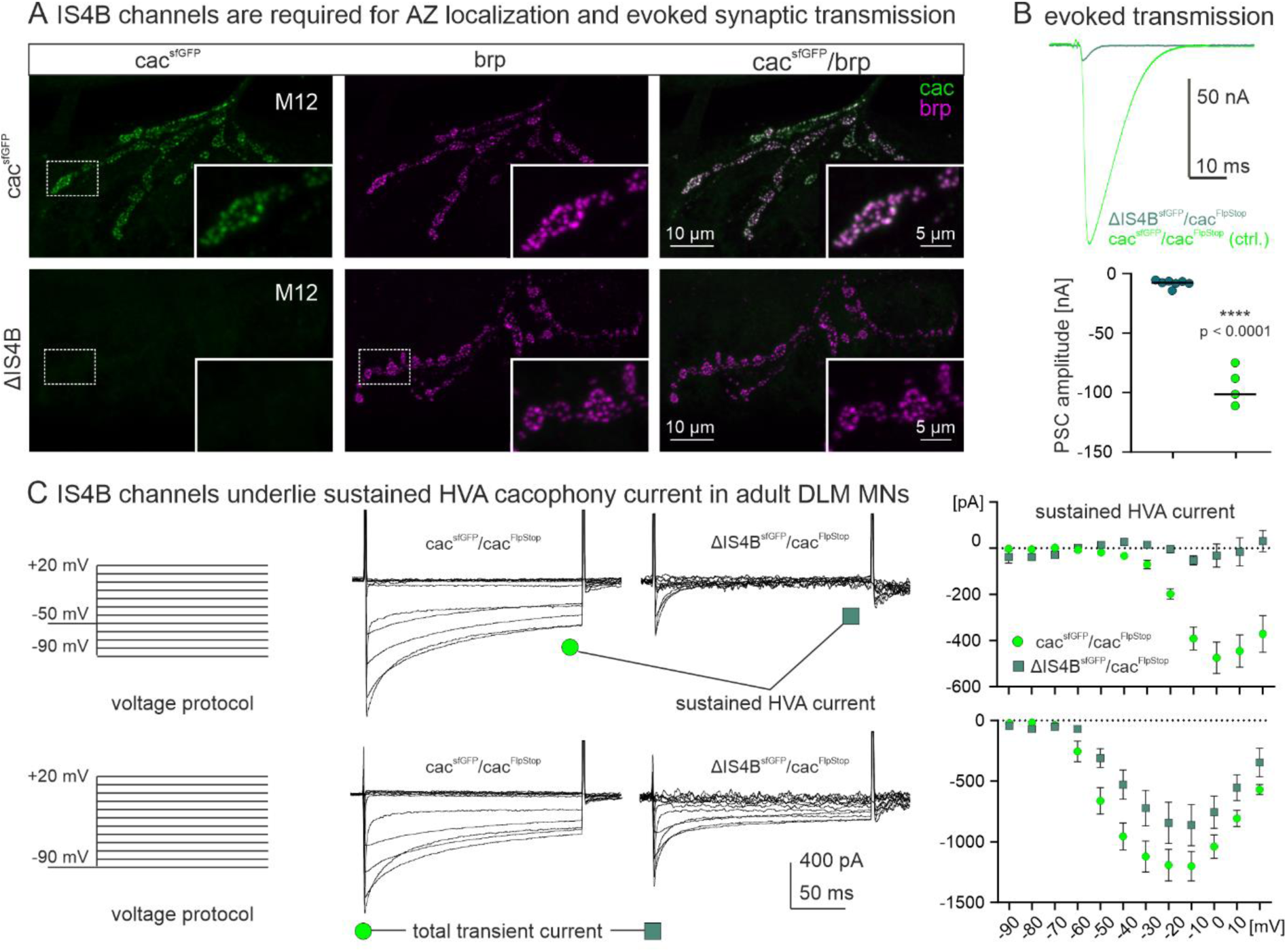
The IS4B exon is required for cacophony localization to AZs and for evoked synaptic transmission. **(A, B)** Since animals homozygous for *IS4B* exon excision are lethal, we created mosaic animals that were heterozygous for cac in most neurons but hemizygous for either cac^sfGFP^ or *ΔIS4B* in motoneurons innervating muscle M12 (see methods, *cac^FlpStop^*). In controls with all cac exons (cac^sfGFP^, green, top row), cac^sfGFP^ colocalizes with brp (magenta) in presynaptic AZs on M12 **(A,** enlargements of area indicated by dotted white rectangle in bottom right corner of each image) and evoked synaptic transmission induces EPSCs of about 100 nA amplitude **(B)**. By contrast, upon deletion of *IS4B* (**A**, bottom row, see also enlargement in bottom right corner) in motoneurons to M12 no cac label is found throughout the motor terminals that are marked by brp (magenta) and evoked synaptic transmission is reduced by more than 90%, Student’s T-test, p < 0.0001 **(B)**. **(C)** Cacophony mediates HVA as well as LVA calcium currents in adult flight motoneurons. All currents are recorded from the somata of adult flight motoneurons in mosaic animals with only one copy of the cac locus in flight motoneurons (see methods). HVA currents (upper traces) are measured by starting from a holding potential of –50 mV (LVA inactivation) followed by step command voltages from –90 mV to +20 mV in 10 mV increments (left, top row), while total cac current (lower traces) is elicited by step commands in 10 mV increments from a holding potential of –90 mV allowing activation of LVA currents (left, bottom row). In GFP-tagged controls (*cac^sfGFP^ / cac^FlpStop^*) this reveals transient and sustained HVA current components (middle, top traces). However, following excision of the *IS4B* exon (*ΔISAB^sfGFP^ / cac^FlpStop^*), the sustained HVA current is absent. By contrast, upon excision of IS4B, the total cac current that also contains cac LVA currents is only partially decreased (middle, bottom traces). Current-voltage (IV) relation of sustained HVA and for total cac current for controls with all cac exons (cac^sfGFP^, green circles, n = 7) and following excision of *IS4B* (*ΔIS4B^sfGFP^*, dark green squares, n = 4).

Previous studies showed that presynaptic cac channels are required for normal evoked synaptic transmission at the Drosophila NMJ (Kawasaki et al. 2002). Reduced cac label in presynaptic AZs goes along with reduced synaptic transmission (Kittel et al. 2006), whereas increased cac channel numbers during presynaptic homeostatic plasticity are effective to increase mean quantal content (Ghelani et al. 2023; Medeiros et al. 2024). If cac presynaptic AZ localization requires *IS4B* but not the *IS4A* exon, removal of *IS4B* but not *IS4A* should impair evoked synaptic transmission. Two electrode voltage clamp recordings from larval muscle 6 in abdominal segment 3 indicate that excitatory postsynaptic currents (EPSCs) as evoked by single presynaptic action potentials are similar in wildtype (Canton S, Figs. 2E, F, blue) and in animals with sfGFP tagged cac channels (Figs. 2E, F, green). Upon removal of *IS4A*, EPSC amplitude is slightly but not significantly increased (Fig. 2E, F, *ΔIS4A*, red). In transheterozygous animals (*ΔIS4B/ΔIS4A*) with removal of *IS4A* on one chromosome and *IS4B* on the other, EPSC amplitude is significantly reduced (Figs. 2E, F, black). Spontaneous synaptic vesicle (SV) release as characterized by the amplitude (Fig. 2G) and the frequency (Fig. 2H) of miniature excitatory postsynaptic currents (mEPSCs) is not affected in *ΔIS4B/ΔIS4A* transheterozygotes. Together, these data indicate that *IS4A* is neither required for AZ localization of channels, nor for evoked synaptic transmission, whereas *IS4B* is required for normal AZ localization and synaptic transmission.

A limitation of these experiments is that homozygous removal of *IS4B* is embryonic lethal, so that presynaptic terminals that are devoid of cac^IS4B^ channels can only be studied in mosaic animals that are heterozygous for *IS4B* excision at most synapses but hemizygous *ΔIS4B* at few synapses of interest. Employing the FlpStop method (see methods, Fisher et al. 2017), mosaic animals with motoneurons to muscle 12 (M12) that are *ΔIS4B/*cac*^null^* can be produced in otherwise heterozygous animals (*ΔIS4B*; Fig. 3A, bottom row). In control animals, cac^sfGFP^ channels localize to brp positive AZs in motoneuron axon terminals on M12 (Fig. 3A, top row and selective enlargement). By contrast, cac^IS4A^ do not localize to brp positive AZs in motoneuron axon terminals on M12 (Fig. 3A, bottom row and selective enlargement). Therefore, *IS4B* is indeed required for cac AZ localization in excitatory glutamatergic axon terminals at the Drosophila NMJ. Consequently, evoked synaptic transmission is nearly abolished in motoneuron terminals that lack IS4B (Fig. 3B). Please note that even upon *IS4B* excision, and thus without any cac channels in AZs, the intensity of brp puncta remained unchanged as compared to control (p = 0.81; Mann Whitney U-test). We conclude that *IS4B* is required for evoked synaptic transmission from chemical presynaptic terminals, whereas *IS4A* has no essential function for action potential induced neurotransmitter release from presynaptic terminals.

### The IS4B exon is required for sustained HVA cac mediated calcium current

Technical constraints prohibit a voltage clamp characterization of IS4B containing cac channels at the larval motoneuron presynaptic terminal, and the somatodendritic calcium current in larval motoneurons is mediated by the Drosophila *Ca*_v_*1* homolog *DmCa1D* (Worrell, Levine 2008; Kadas et al. 2017). However, Drosophila cac currents have previously been shown in somatodendritic voltage clamp recordings from pupal and adult motoneuron somata (Ryglewski et al. 2012; 2014a; b). Although these recordings are not representative for cac currents at larval presynaptic terminals, they show that Drosophila cac channels can in principle contribute to transient and sustained as well as high (HVA) and low (LVA) voltage activated currents (Ryglewski et al. 2012). Given that mutually exclusive splicing at the *IS4* site affects the voltage sensor (Fig. 1B) and that only the *IS4B* exon is required for evoked synaptic transmission, we next tested whether *IS4B* makes a significant contribution to a specific sub-type of cac mediated calcium current. The comparison of voltage clamp recordings from adult flight motoneuron somata in animals with full cac isoform diversity and mosaic animals with *IS4B* excision in only these motoneurons (see methods) shows that *IS4B* is required for sustained HVA cac current. Following electrical inactivation of LVA currents with prepulses to –50 mV (Fig. 3C, top left), HVA with an activation voltage of ∼-30 mV shows a sustained component that is reliably recorded with full cac isoform diversity (Fig. 3C, top left current traces and top IV diagram) but the sustained HVA component is absent in *ΔIS4B* motoneurons (Fig. 3C, top right current traces and top IV diagram). By contrast, total cac current (bottom traces) that contains LVA and HVA components is not abolished but only reduced by the excision of *IS4B* (Fig. 3C, bottom). Somatodendritic calcium current as recorded in neurons without the IS4B exon prove that cac^IS4A^ channel isoforms are expressed and functional (Fig. 3C) although cac^IS4A^ isoforms are not present in presynaptic terminals at the NMJ (Figs. 2C, 3A). Therefore, the *IS4B* exon is vital for evoked synaptic transmission and promotes sustained HVA calcium current, whereas the *IS4A* exon is not sufficient for evoked synaptic transmission but mediates somatodendritic cacophony current in adult flight motoneurons.

### Alternative splicing at the I-II site does not affect active zone localization but channel number and release probability

Immunohistochemical triple label at the larval Drosophila NMJ tests whether either one of the mutually exclusive exons at the *I-II* locus is required for correct presynaptic cac localization. Motoneuron axon terminals on larval muscles 6 and 7 (M6/7) are labeled with HRP (Fig. 4A-C, right column, blue), sfGFP tagged cac channels by α-GFP immunolabel (Fig. 4A-C, left column, green), and AZs by α-brp immunocytochemistry (Fig. 4A-C, second column, magenta). For a better visualization of brp and cac puncta, selective enlargements are shown for each genotype in Figs. 4Ai-Ci. In controls, sfGFP tagged cac channels with full isoform diversity co-localize with brp in presynaptic AZs (Figs. 4A, Ai) as also shown above (Figs. 2A, Ai). The same is the case for both mutually exclusive variants at the *I-II* locus. Cac channels that contain I-IIB but lack I-IIA (*ΔI-IIA*, Figs. 4B, Bi), and *vice versa*, cac channels that contain I-IIA but lack I-IIB (*ΔI-IIB*, Fig. 4C, Ci) show AZ localization (see overlays of brp and cac^sfGFP^ in Figs. 4B, Bi, C, Ci, third column). For all cac isoforms that are targeted to presynaptic AZs, quantification reveals similar Pearson’s correlation coefficients of ∼0.65 (Fig. 4D), which corresponds to previous reports (Krick et al. 2021). Moreover, the Manders 1 and 2 co-localization coefficients are similar for control, cac^IS4B^, and both I-II locus isoforms cac^I-IIB^ and cac^I-IIA^ (Figs. 4E, F). In sum, alternative splicing at the *I-II* site does not affect cac expression in presynaptic AZs.

**Figure 4.**
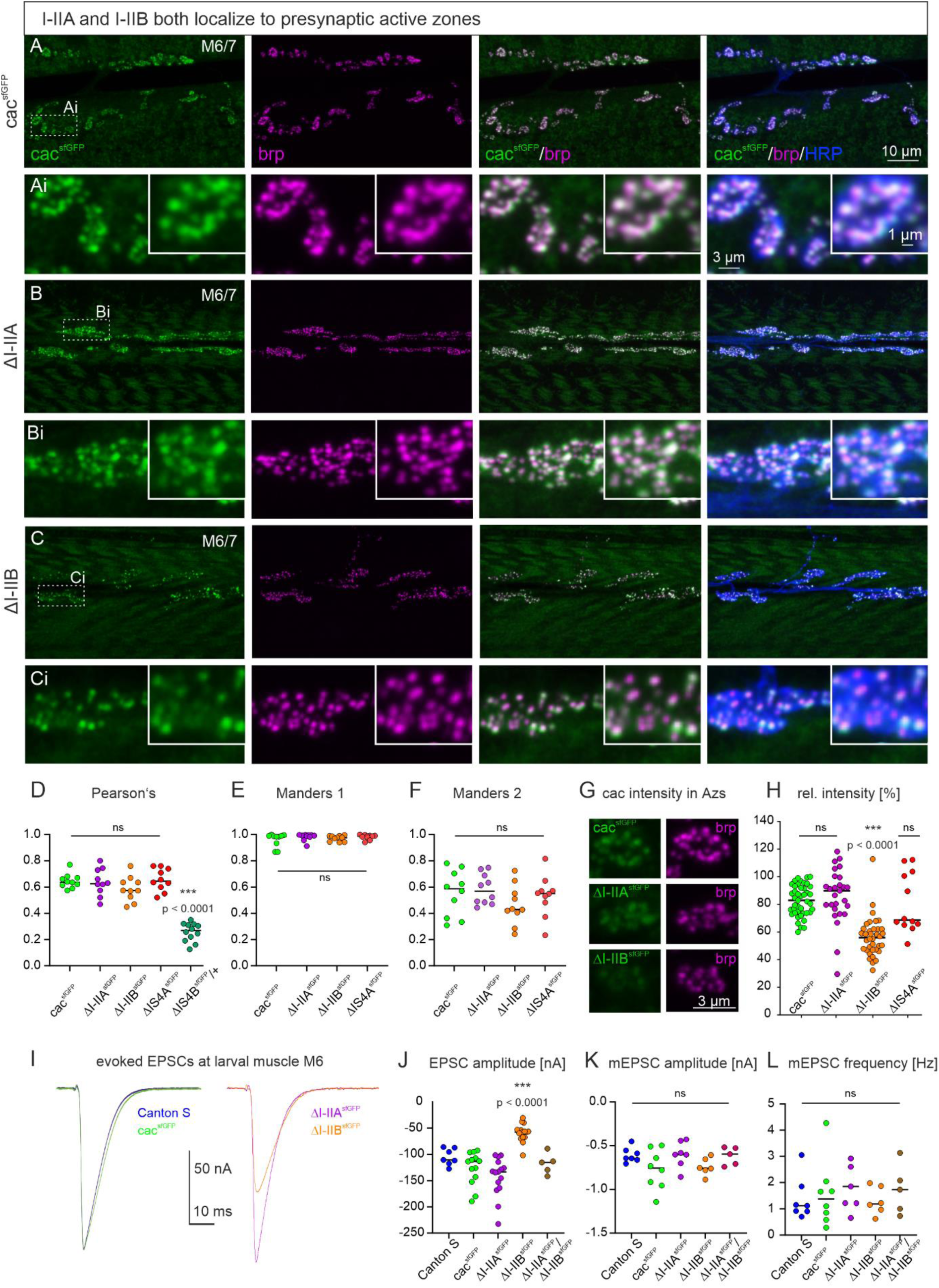
Excisions at the I-II exon do not affect AZ cacophony localization but can alter cac^sfGFP^ label intensity in AZs and EPSC amplitude. **(A-C)** Representative confocal projection views of triple labels for GFP tagged cac channels (green), the AZ marker brp (magenta), and HRP to label axonal membrane (blue) with enlargements that are indicated by white dotted rectangles in A-C (Ai-Ci). Label in control animals with all cac exons (cac^sfGFP^, top two rows, **A, Ai**), with selective excision of either the alternative exon *I-IIA* (*ΔI-IIA^sfGFP^*, middle two rows, **B, Bi**), or the alternative exon *I-IIB* (*ΔI-IIB^sfGFP^*, bottom two rows, **C, Ci**). The gross morphology of the neuromuscular junctions (muscle fibers, bouton numbers and sizes, AZ numbers) was similar in all three genotypes (not shown). Excision of the *I-IIA* exon does neither impact cac AZ localization nor labeling intensity **(B, Bi, H)**. Excision of *I-IIB* does not impact cac AZ localization but labeling intensity seems lower **(C, Ci, H)**. **(D-F)** Quantification of cac co-localization with the AZ marker brp yields a similar Pearson’s colocalization coefficient **(D)** as well as similar Manders 1 **(E)** and Manders 2 **(F)** coefficients for controls (green) and both exon-out variants of the *I-II* locus (ΔI-IIA purple, ΔI-IIB orange) as well as for *ΔIS4A* (red) but not for *ΔIS4B* (dark green, ANOVA with Dunnett’s post hoc test, p < 0.0001). **(G)** ΔI-IIB^sfGFP^ shows fainter immunofluorescence signals in the AZ as compared to control (cac^sfGFP^) and *ΔI-IIA^sfGFP^*. **(H)** Quantification confirms a significant reduction in *ΔI-IIB^sfGFP^* labeling intensity (Kruskal Wallis ANOVA with Dunn’s post hoc test, p < 0.0001) and no differences between *ΔI-IIA^sfGFP^* and control (p > 0.99). **(I)** Evoked synaptic transmission as recorded in TEVC from muscle fiber 6 upon extracellular stimulation of the motor nerve. Excitatory postsynaptic currents (EPSCs) are of similar shape and amplitude for CS control (blue) and animals with GFP-tagged cac channels (cac^sfGFP^, green, p=0.34, two sided Tukey’s multiple comparison test). Excision of *I-IIA* (*ΔI-IIA^sfGFP^*, purple) has no effect on evoked release amplitude (p=0.52, two sided Tukey’s multiple comparison test), but excision of *I-IIB* (*ΔI-IIB^sfGFP^*, orange) reduces evoked release significantly (p < 0.0001). **(J)** Quantification of EPSC amplitude reveals a highly significant reduction in *ΔI-IIB^sfGFP^* (orange) as compared cac^sfGFP^ controls (green), but neither animals with excision of the *I-IIA* exon (purple), nor transheterozygous animals with excision of *I-IIA* on one and *I-IIB* on the chromosome (brown) show differences to control (p=0.97, two sided Tukey’s multiple comparison test). **(K)** Quantal size (mEPSC amplitude) and spontaneous release frequency **(L)** show no significant difference among genotypes.

However, removal of the *I-IIB* exon reduces the intensity of cac immunolabel in AZs as compared to control or *ΔI-IIA* highly significantly by ∼50% (Figs. 4G, H). These data indicate that *I-IIB* excision may result in fewer cac channels per presynaptic AZ. We employ two electrode voltage clamp recordings from the postsynaptic cell (M6) to test for the resulting consequences on synaptic transmission (Figs. 4I-L). The amplitude of the postsynaptic current (EPSC) as evoked by an action potential in the presynaptic motor axon is similar in Canton S with untagged cac and animals expressing cac^sfGFP^ channels (Figs. 4I, J, see also above, Figs. 2E, F). Removal of the *I-IIA* exon (*ΔI-IIA*, Fig. 4I, purple trace) does not result in significant differences of EPSC amplitude as compared to tagged or untagged controls (Figs. 4I, J). By contrast, removal of the *I-IIB* exon (*ΔI-IIB*, Fig. 4I, orange trace) results in a reduction in EPSC amplitude by ∼50% (Fig. 4J), which matches the reduced channel immunofluorescence signal in AZs (Fig. 4H) upon removal of the *I-IIB* exon. By contrast, neither removal of the *I-IIA* exon nor removal of the *IS4A* exon reduces tagged cac channel intensity in AZs (Fig. 4H), and neither of these manipulations affect EPSC amplitude (Figs. 2F, 4J). The differences or similarities between genotypes in EPSC amplitude also hold when analyzing EPSC charge instead of amplitude (significantly smaller EPSC charge in *ΔI-IIB*, p < 0.0001, ANOVA with Dunnet post hoc comparison). In contrast to different cac puncta intensities in I-IIA versus I-IIB containing cac channels the intensity of brp puncta remains control like across the exon excision mutants (Kruskal Wallis ANOVA, p = 0.1, data not shown).

It seems unlikely that presynaptic cac channel isoform type affects glutamate receptor types or numbers, because the amplitude of spontaneous miniature postsynaptic currents (mEPSCs, Fig. 4K) and the labeling intensity of postsynaptic GluRIIA receptors are not significantly different between controls, I-IIA, and I-IIB junctions (see suppl. Fig. 2, p = 0.48, ordinary one-way ANOVA, mean and SD intensity values are 61.0 ± 6.9 (control), 55.8 ± 8.5 (*ΔI-IIA*), 61.1 ± 17.3 (*ΔI-IIB*)). However, we cannot exclude altered GluRIIB numbers and have not quantified GluR receptor field sizes. Similarly, the frequency of miniature postsynaptic currents (mEPSCs) remains unaltered (Fig. 4L). Since mEPSC frequency has been related to RRP size at some synapses (Pan et al., 2009; Ralowicz et al., 2024), this indicates unaltered RRP size upon *I-IIB* excision, but we have not directly measured RRP size. In sum, these data show that neither exon at the *I-II* locus is required for cac localization to AZs, but *I-IIB* is required for normal evoked synaptic transmission amplitude. A reduced amplitude of evoked synaptic transmission along with less intensive presynaptic cac immunolabel in *ΔI-IIB* animals (Figs. 4H, J) is indicative for fewer calcium channels in AZs. Alternatively, the nanoscale localization of cac could be affected by alternative splicing.

We assess the latter by dual color STED microscopy of the AZ marker brp and cac^sfGFP^ in different exon excision mutants (Figs. 5A-E). Collecting STED image stacks of synaptic boutons on muscle 6 (Fig. 5A) reveals numerous AZs in various spatial orientations relative to the focal plane (Fig. 5A-C). In a strict top view (Fig. 5C1) the orientation of the synapse is planar so that the central cac cluster (magenta) is surrounded by four brp puncta (green) all in the same plane. If the AZ lies tilted relative to the focal plane, the same arrangement is viewed from different angles (side views, Figs. 5C2-6). The localization of cac relative to brp in different exon-out variants (Figs. 5D, E) is quantified by measuring the distance between the center of the cac cluster and the center of the nearest brp punctum in top and side views within the same optical sections (see methods). There is no difference in the cac to brp distance between different views/synapse orientation in relation to the focal plane. In all cac exon-out variants that are expressed in AZs (control, *ΔIS4A*, *ΔI-IIA* and *ΔI-IIB* but not *ΔIS4B*) the median distance ranges between 103 and 109 nm and reveals no significant differences (Kruskal-Wallis test, p = 0.62) between control (median distance, 106.1 nm) and any of the exon-out variants shown *(ΔIS4A*, *ΔI-IIA* and *ΔI-IIB*, Fig. 5E). Alternative splicing at the *I-II* locus does therefore neither affect targeting of cac to AZs (Figs. 4A-F), nor cac localization within the brp scaffold of the AZ (Figs. 5A-E).

**Figure 5.**
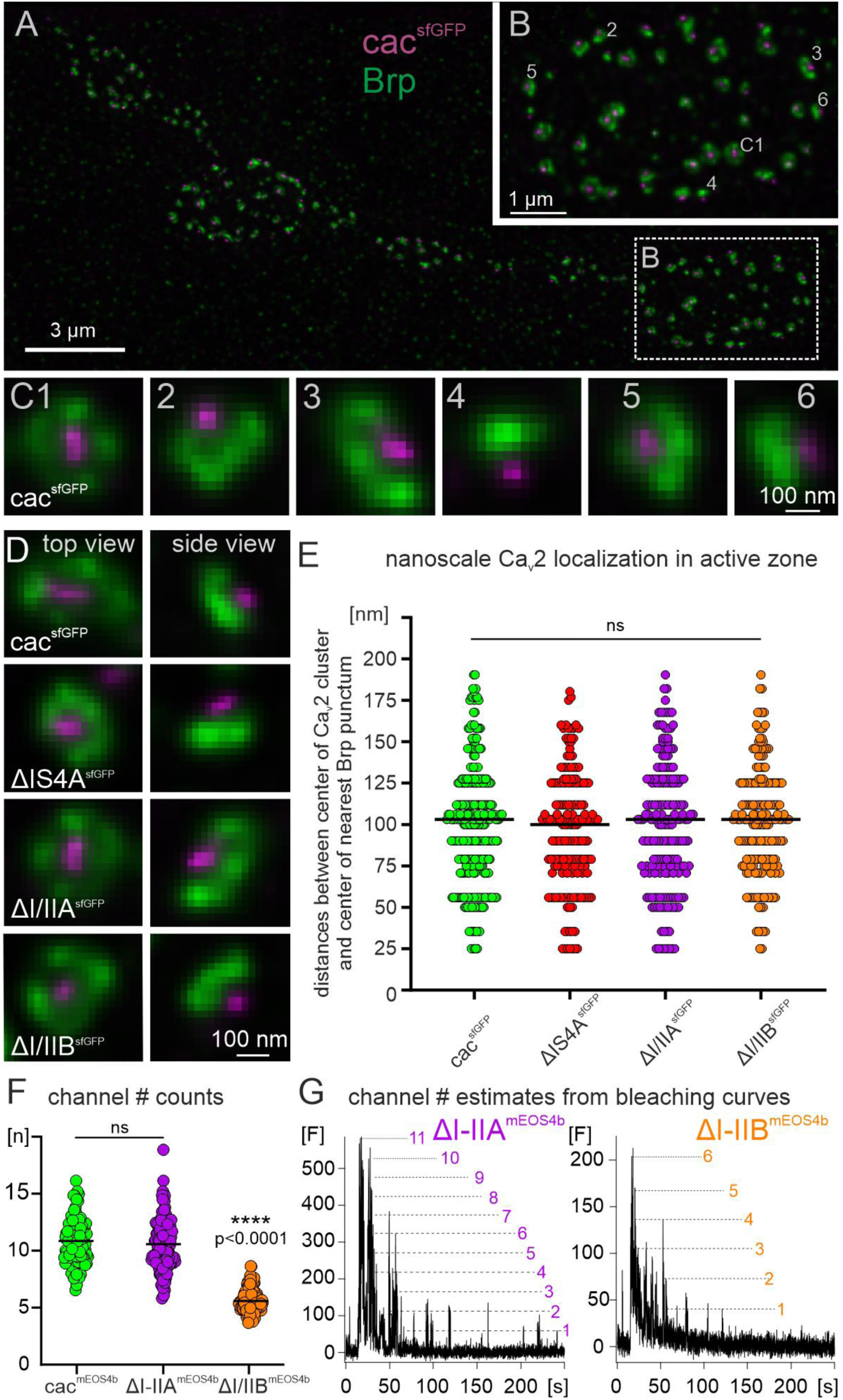
Dual color STED imaging reveals equal nanoscale channel localization in AZs of cacophony for all exon-out variants, and live sptPALM imaging reduced channel numbers in AZs for ΔI-IIB. **(A)** Representative intensity projection image of the AZ marker bruchpilot (labeled with anti-brp, green) and cac clusters (cac^sfGFP^ labeled with anti-GFP, magenta) as imaged with dual color STED at motoneuron axon terminal boutons on larval muscle M6. The dotted white box demarks one bouton that is enlarged in **(B)**. Each cac cluster (magenta) is in close spatial proximity to the AZ marker brp (green) and depending on AZ orientation, cac and brp is viewed from different angles. Top views (= planar views, see C1 in B and in selective enlargement) show 4 brp puncta that symmetrically surround the central cac cluster. Viewing AZs at the edge of the bouton shows the cac cluster facing to the outside and the brp puncta in close proximity (see 2-6). **(C)** Selective enlargements of each AZ that is numbered in B. **(D)** Top views (left column) and side views (right column) of the cac/brp arrangement in AZs in controls with cac^sfGFP^ (top row), with excision of exon *IS4A* (*ΔISA4^sfGFP^*, second row), with excision of exon *I/IIA* (*ΔI/IIA^sfGFP^*, third row), and with excision of exon *I/IIB* (*ΔI/IIB*^sfGFP^, bottom row). **(E)** Quantification of the distances between the center of each cac punctum to the nearest brp punctum in the same focal plane. **(F-G)** Live sptPALM imaging of mEOS4b tagged cac channels from AZs of MN terminals on muscle 6 in controls with full isoform diversity (cac^mEOS4b,^ green) and following the removal of either *I-IIA* (*ΔI-IIA^mEOS4b^*, purple) or *I-IIB* (*ΔI-IIB^mEOS4b^*, orange). **(F)** Quantification of channel numbers from bleaching curves **(G)** reveals ∼ 9-11 cacophony channels per AZ for tagged controls, which matches previous reports (Ghelani et al. 2023). Counts for *ΔI-IIA* reveal no significant differences (Kruskal-Wallis test with Dunn’s posthoc comparison, p=0.94), but cac number in AZs is reduced by ∼50 % in *ΔI-IIB* (p<0.0001). **(G)** Bleaching curves of single AZs were illuminated after ∼10 s and then imaged under constant illumination for another 240 s. Discrete bleaching steps (dotted lines) indicate the bleaching of single cac^mEOS4b^ channels. Comparing the amplitudes of single events and their integer multiples (dotted lines) to the maximum fluorescence at illumination start allows estimates of the total channel number per AZ.

This leaves changes in cac properties or channel number in AZs as plausible causes for the reduction in evoked synaptic transmission upon removal of the *I-IIB* exon (Figs. 4I, J). As previously reported, channel number in presynaptic AZs can be estimated by live sptPALM imaging of mEOS4b tagged cac channels at the NMJ (cac^mEOS4b^, Ghelani et al. 2023). To estimate channel number in AZs of axon terminals on larval muscle M6 in controls, *ΔI-IIA*, and *ΔI-IIB* (Fig. 5F), the bleaching behavior of cac^mEOS4b^ signals in individual AZs is imaged during steady illumination. Discrete bleaching steps (Fig. 5G, dotted lines) indicate bleaching events of individual cac^mEOS4b^ molecules, and thus the fluorescence intensity of a single cac^mEOS4b^ channel. Larger channel numbers produce integer multiple fluorescence intensity amplitudes. Dividing the full fluorescence amplitude that is measured at the illumination onset of all channels in the AZ by the fluorescence intensity from a single channel yields total channel number per AZ. Quantification from three animals per genotype with at least 30 AZs per animal confirms a previous study (Ghelani et al. 2023) showing that control animals with full cac channel isoform diversity express ∼10 cac channels per AZ (Fig. 5F, green, 10.8 ± 2 channels). Removing the alternative exon *I-IIA* does not affect channel number per AZ (Fig. 5F, purple, 10.5 ± 2 channels), but excision of *I-IIB* reduces channel number to ∼50 % (Fig. 5F, orange, 5.6 ± 1 channels). A ∼50% reduction in channel number counts in AZs (Fig. 5F) is in line with ∼50 % reduction in cac immunofluorescence in AZs (Fig. 4H) and evoked synaptic transmission (Figs. 4I, J) upon excision of *I-IIB*. In sum, these data indicate that the reduction of evoked synaptic transmission amplitude in *ΔI-IIB* is a consequence of reduced channel number.

### Functional consequences of I-II site alternative splicing during repetitive firing

In addition to reducing the numbers of SVs that are released upon one presynaptic action potential (quantal content), removal of the *I-IIB* exon has significant effects on synaptic transmission during repetitive stimulation and on synaptic plasticity. First, the paired pulse ratio (PPR) is affected. In 0.5 mM calcium, controls with sfGFP-tagged cac channels show slight paired pulse (PP) depression at interpulse interval (IPI) durations below 20 ms (Fig. 6A). Similarly, upon removal of the *I-IIA* exon (*ΔI-IIA^sfGFP^*) PP depression is observed for IPIs below 20 ms (Fig. 6B). By contrast, upon removal of the *I-IIB* exon, some animals show PP depression and others PP facilitation, so that the average PPR is close to 1 for all IPIs, but the variance is large, in particular for short IPIs (Fig. 6C). To test whether increased PPRs and increased PPR variance can be fully explained by reduced channel numbers upon excision of *I-IIB*, or whether additional factors were required to explain these findings, we increased the external calcium concentration to 1.8 mM so that the first pulse amplitude in *ΔI-IIB* animals matched that of control animals in 0.5 mM calcium. This fully rescued the effect of *I-IIB* excision. In fact, mean and variance of PPRs as recorded from *ΔI-IIB* in 1.8 mM external calcium were similar to the PPRs as observed in controls and in *ΔI-IIA* in 0.5 mM external calcium across all interpulse intervals (Fig. 6D). Considering the variance at 0.5 mM external calcium, for control and for excision of *I-IIA* the coefficient of variation is ∼5-10 % across IPIs whereas upon excision of *I-IIB* it reaches 15-20% for short IPIs (Fig. 6E). However, increasing external calcium to 1.8 mM in recordings in *ΔI-IIB* animals increases the first EPSC amplitude to that observed in controls in 0.5 mM external calcium (Fig. 6D) and eliminates altered PPRs as well as increased variance (Figs. 6D, E).

**Figure 6.**
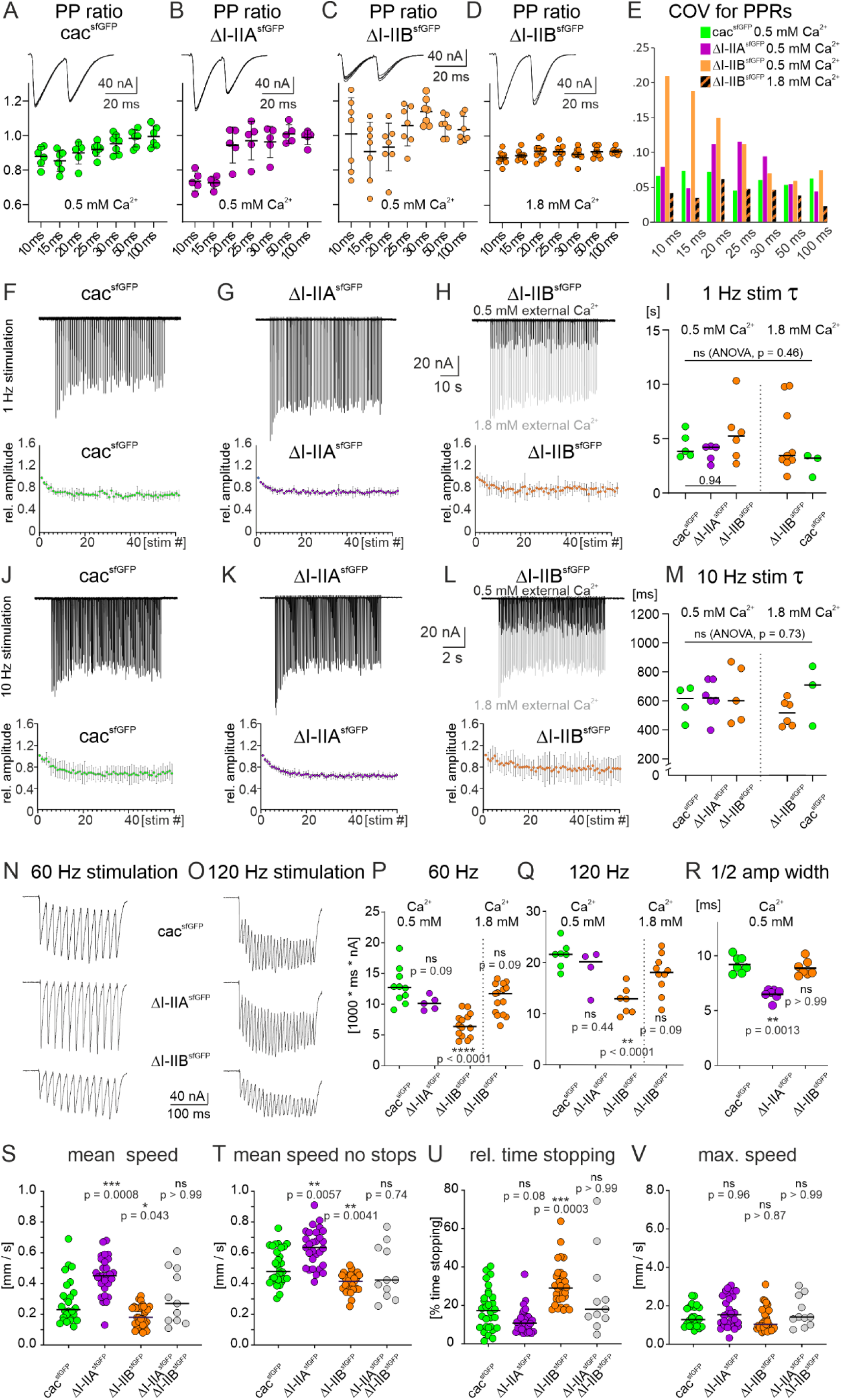
Alternative splicing in the I-II linker affects short term plasticity due to decreased calcium influx and motor behavior. **(A-D)** Paired pulse ratio (PPR, ratio of second EPSC amplitude divided by first EPSC amplitude) as measured in 0.5 mM external calcium (**A-C**) or in 1.8 mM external calcium (**D**) at different interpulse intervals (IPIs ranging from 10 ms to 100 ms) in control animals (cac^sfGFP^, **A**), in animals with removal of *I-IIA* (*ΔI-IIA^sfGFP^*, **B**), and in animals with removal of *I-IIB* (*ΔI-IIB^sfGFP^*) either in 0.5 mM calcium (**C**) or 1.8. mM calcium (**D**). The large variance in PPR upon excision of I-IIB (**C**) is rendered control-like if the first EPSC amplitude of the twin pulse is adjusted to 0.5 mM external calcium control level (**D**; comp. with **A**). This is also reflected in the coefficient of variation (COV) for PPRs (**E**, 0.5 mM calcium: cac^sfGFP^ green, *ΔI-IIA* purple, *ΔI-IIB* orange; 1.8 mM calcium: *ΔI-IIB* orange/black pattern). **(F-I)** Synaptic depression as measured in 0.5 mM external calcium in response to stimulus trains of 1 minute duration at 1 Hz frequency for animals with GFP-tagged cac (cac^sfGFP^, **F**), following removal of *I-IIA* (*ΔI-IIA^sfGFP^*, **G**), and with removal of *I-IIB* (*ΔI-IIB^sfGFP^*, **H**). The top traces show representative TEVC recordings from the postsynaptic muscle cell, and the diagrams mean values (n=5 for F and G, N=6 for H, error bars are SD). (**H**) The light gray trace shows *ΔI-IIB* in 1.8 mM external calcium, while the black trace shows *ΔI-IIB* in 0.5 mM calcium. For all 3 genotypes, depression reaches steady state at ∼ 80 % of the original EPSC amplitude, but upon excision of *I-IIB* it is more variable **(H)**. Depression time courses do not differ between genotypes but are more variable in *ΔI-IIB,* independent of external calcium concentration (**H, I**). **(J-M)** Synaptic depression in response to stimulus trains at 10 Hz frequency for animals with GFP-tagged cac (cac^sfGFP^, **J**), following removal of *I-IIA* (*ΔI-IIA^sfGFP^*, **K**), and with removal of *I-IIB* (*ΔI-IIB^sfGFP^*, **L**: black in 0.5 mM calcium, gray trace in 1.8 mM calcium). Again, depression is most variable between animals upon excision of *I-IIB* **(L, M)** but time courses do not differ between genotypes. However, time course variation decreases in 1.8 mM calcium in animals with excision of *I-IIB* (**M**). Motoneuron stimulation at 60 **(N)** or 120 Hz **(O)** frequency, both for durations of 200 ms in animals with GFP-tagged cac (cac^sfGFP^, top traces), following removal of *I-IIA* (*ΔI-IIA^sfGFP^*, middle traces), and with removal of *I-IIB* (*ΔI-IIB^sfGFP^*, bottom traces). To compare charge transfer across the NMJ during high frequency bursts the total EPSC area below baseline (prior to stimulation) was measured during each 200 ms burst and plotted for each genotype for 60 Hz stimulation in **(P)** and for 120 Hz stimulation in **(Q).** Decreased charge transfer in animals with excision of the *I-IIB* exon is rescued to control level if external calcium is increased to 1.8 mM so that the first EPSP matches control amplitude in 0.5 mM external calcium (**P, Q,** far right data points *ΔI-IIB* in 1.8 m calcium). **(R)** shows single evoked EPSC half amplitude width. (**S-V**) show different measurements during larval crawling for control animals with GFP-tagged cac (cac^sfGFP^), removal of *I-IIA* (*ΔI-IIA^sfGFP^*), removal of *I-IIB* (*ΔI-IIB^sfGFP^*), and in transheterozygous animals with removal of *I-IIA* on one and removal of *I-IIB* on the other chromosome (*ΔI-IIA^sfGFP^/ΔI-IIB^sfGFP^*). The measured parameters are mean speed during 10 minutes of crawling **(S)**, mean speed without any stops **(T)**, the relative time spent stopping **(U)** and the maximum speed reached **(V)**. In all diagrams each dot demarks a measurement from a different animal and horizontal bars the medians. For statistics, non-parametric Kruskal Wallis ANOVA with planned Dunn’s posthoc comparison to control was conducted.

We next tested whether the time course of synaptic depression during low frequency repetitive stimulation at 1 Hz is affected upon removal of either *I-II* exon. In 0.5 mM external calcium, repetitive stimulation of the NMJ to muscle M6/7 at 1 Hz frequency for 1 minute causes synaptic depression that reaches steady state at about 80 % of the initial transmission amplitude with a time constant of about 5 s (Krick et al. 2021). The same is observed for cac^sfGFP^ controls (Figs. 6F, I) and upon removal of *I-IIA* (Figs. 6G, I). Removal of *I-IIB* does not affect the magnitude and time course of depression at 1 Hz stimulation (Fig. 6H), but it increases the variability of the time constant until steady state depression is reached (Fig. 6I). Again, we triturated the external calcium concentration (1.8 mM) so that the first EPSC amplitude in the train in Δ*I-IIB* matched the first one in controls recorded in 0.5 mM external calcium (gray trace in Fig. 6H is *ΔI-IIB* in 1.8 mM calcium and black trace in 0.5 mM calcium). This did neither affect the mean time constant of depression nor the larger variance observed in *ΔI-IIB* (Fig. 6I), indicating that these parameters do not depend on calcium influx into AZs. At 10 Hz stimulation neither removal of *I-IIA* nor of *I-IIB* affects the time course or amplitude of synaptic depression (Figs. 6J-M). Increasing the external calcium concentration (1.8 mM) so that the first EPSC amplitude in the train in *ΔI-IIB* matches the first one in controls recorded in 0.5 mM external calcium (gray trace in Fig. 6L is *ΔI-IIB* in 1.8 mM calcium and black trace in 0.5 mM calcium) does not affect magnitude or time course of depression (Fig. 6M). Similarly, increasing external calcium to 1.8 mM in controls does not affect the time constant of synaptic depression, in line with the findings at 1 Hz that these parameters do not depend on the amount of calcium influx into AZs.

Taken together removal of *I-IIA* does not alter channel number and does not affect PPRs or the time course of synaptic depression. By contrast, removal of *I-IIB* reduces channel number. This in turn is the cause for altered PPRs at short interpulse intervals (Figs. 6C, E) and can be rescued by increasing external calcium. Neither excision of *I-IIA*, nor of *I-IIB* has significant effects on the magnitude or time course of synaptic depression at low stimulation frequencies.

However, motoneuron firing frequencies as previously measured during tethered crawling (Kadas et al. 2017) reach ∼120 Hz during bursts of about 200 ms duration. Given that the main function of excitatory synaptic transmission to larval muscles is locomotion, it seems important to test stimulation protocols that reflect behaviorally relevant motoneuron firing patterns. Applying motoneuron stimulation for 200 ms duration at either 60 Hz (Fig. 6N) or 120 Hz frequency (Fig. 6O) in 0.5 mM external calcium reveals summation of the EPSCs in control animals with GFP-tagged cac channels (Figs. 6N, O, top traces) as well as in animals without the *I-IIB* exon (*ΔI-IIB*, Figs. 6N, O, bottom traces). By contrast, upon excision of *I-IIA*, summation of EPSCs is absent at 60 Hz but present at 120 Hz stimulation frequency (*ΔI-IIA*, Figs. 6N, O, middle traces). The reason why EPSP summation occurs only at very high motoneuron firing frequencies (120 Hz) in *ΔI-IIA*, but already at 60 Hz in controls and in *ΔI-IIB* animals is a significantly smaller EPSP half-width upon excision of the *I-IIA* exon (Fig. 6R). This might be an indication for an effect of *I-IIA* exon excision on cac channel biophysical properties (see discussion). At both high frequency stimulation protocols (60 and 120 Hz) charge transfer is significantly reduced upon excision of *I-IIB* (Figs. 6P, Q). We measure charge transfer as the total area under the postsynaptic response traces during 200 ms long stimulation bursts at either 60 Hz (Figs. 6N, P) or 120 Hz (Figs. 6O, Q). To test whether this is caused by reduced calcium influx during an EPSC in *ΔI-IIB* (Figs. 6N, O), we again increased the external calcium concentration to 1.8 mM so that the first EPSC of the train matches the first EPSC amplitude observed in controls in 0.5 mM external calcium (see Figs. 6N, O, bottom gray overlay traces). This increased median total charge in *ΔI-IIB* to control values for both 60 Hz and 120 Hz stimulation (Figs. 6P, Q), thus indicating that the reduced charge transfer in *ΔI-IIB* at 0.5 mM external calcium can be explained by reduced presynaptic calcium channel number and concomitant reduced calcium influx.

Although crawling speed of Drosophila larvae is mainly adjusted by varying the duration between subsequent peristaltic waves of motoneuron bursting (peristaltic wave period duration, Liu et al. 2023), differences in cac mediated neuromuscular synaptic transmission may also play an important role. In accordance with reduced charge transfer to the postsynaptic muscle cell upon excision of the *I-IIB* exon (Figs. 6N-R), mean crawling speed is significantly reduced in *ΔI-IIB* animals (Fig. 6S). The mean crawling speed (including stops) of cac^sfGFP^ control larvae is roughly 0.2 mm per second and not significantly affected in transheterozygous *ΔI-IIA* over *ΔI-IIB* larvae that contain full cac isoform diversity (Figs. 6T, U). Removal of *I-IIB* (*ΔI-IIB*) causes a significant decrease in mean locomotion speed (Fig. 6S, orange) whereas removal of *I-IIA* significantly increases speed (*ΔI-IIA*, Fig. 6S, purple). Alterations in locomotion speed upon alternative cac exon removal could either be caused by changes in mean ground speed (mean speed excluding stops), or by changes in the duration of stopping, or by both. Mean ground speed in controls is about 0.5 mm per second (Fig. 6T), which is slightly slower but within the range previously reported (0.65-0.8 mm per second, Wang et al. 1997; Guo et al. 2016). The decreased net locomotion speed in *ΔI-IIB* larvae is caused by significant reductions in the mean ground speed (Fig. 6T) paired with significant increases in the duration of stopping (Fig. 6U). By contrast, the net locomotion speed increase as observed in *ΔI-IIA* larvae is caused by significant increases in mean ground speed (Fig. 6T) without significant changes in the duration of stopping (Fig. 6U). However, the maximum locomotion speed observed does not differ significantly between controls and any of the test groups (Kruskal Wallis test, p=0.14; Fig. 6V), although *ΔI-IIB* neuromuscular junctions show significantly reduced charge transfer at 120 Hz motoneuron bursting (Fig. 6Q). This may indicate a safety plateau of charge transfer at maximum speed.

### I-IIB is required for presynaptic homeostatic plasticity

Chemical synapses are subject to various forms of short-term (Fioravante, Regehr 2011), Hebbian (Nicoll, Schmitz 2005) and homeostatic plastic adjustments (Turrigiano 2008). The Drosophila larval NMJ has become a prominent model to analyze the mechanisms underlying presynaptic homeostatic potentiation (PHP, Frank et al., 2006; Davis, Müller 2015). PHP is a compensatory increase in the number of SVs that are released upon one presynaptic action potential (quantal content) in response to reduced postsynaptic receptor function. Consequently, EPSC amplitude is restored to its original setpoint despite reduced mEPSC amplitudes. Quantal content can be increased by a larger size of the readily releasable pool (RRP) of SVs, or by elevated release probability (P_r_). A recent study has shown that the induction of PHP at the Drosophila NMJ requires an increase in P_r_ that is mediated by increasing the number of cac channels in the presynaptic AZ (from ∼10 to ∼12, Ghelani et al, 2023; see also Gratz et al. 2019). Our data show that exclusion of *IS4B* impairs cac localization to the AZ (Figs. 2C, Ci and 3A), whereas exclusion of *I-IIB* reduces cac number in the presynaptic AZ (Fig. 5F), raising the question whether homeostatic plasticity is affected by *I-II* exon splicing. Acute pharmacological blockade of postsynaptic glutamate receptors with the bee wolf toxin, philanthotoxin (PhTx), is a well-established means to induce PHP within minutes at the Drosophila larval NMJ (Frank et al. 2006; Davis, Müller 2012; 2015; Gratz et al. 2019; Ghelani et al. 2023). In control animals with GFP-tagged cac channels as well as upon excision of the *I-IIA* exon (normal channel numbers, Fig. 5F), bath application of PhTx reliably reduces the amplitude of spontaneously occurring minis (mEPSCs; data are presented as % of baseline without PhTx within genotype: cac^sfGFP^ control: Figs. 7A, E, green; *ΔI-IIA*: Figs. 7B, E, purple), but evoked EPSC amplitude is not reduced as compared to control (data are presented as % of baseline without PhTx; cac^sfGFP^ control: Figs. 7A, D, green; *ΔI-IIA*: Figs. 7B, D, purple). Normal EPSC amplitudes at reduced mEPSC amplitudes are caused by a compensatory increase in mean quantal content (data are presented as % of baseline without PhTx; Fig. 7F; cac^sfGFP^ control: green; *ΔI-IIA*: purple). By contrast, PHP is not observed upon removal of the *I-IIB* exon (Figs. 7C-F). As in controls and in *ΔI-IIA* animals, PhTx application reduces mEPSC amplitude by ∼0.2 nA as expected (Figs. 7C, E), but in *ΔI-IIB* no compensatory increase in mean quantal content is observed (Fig. 7F), so that EPSC amplitude is not restored to its original setpoint (Fig. 7D). Similarly, in *ΔI-IIB* animals, PHP is also not possible in response to a permanent reduction of glutamate receptor function in *GluRIIA* mutants, which has been named PHP maintenance and is genetically separable from PHP induction (James et al. 2019). In *GluRIIA* mutants, compensatory upregulation of mean quantal content is observed in animals with GFP-tagged cac and upon excision of *I-IIA*, but not following excision of the *I-IIB* exon (Fig. 7G). Therefore, the lack of the *I-IIB* exon causes fewer cac calcium channels in the AZ and impairs both PHP initiation and maintenance.

**Figure 7.**
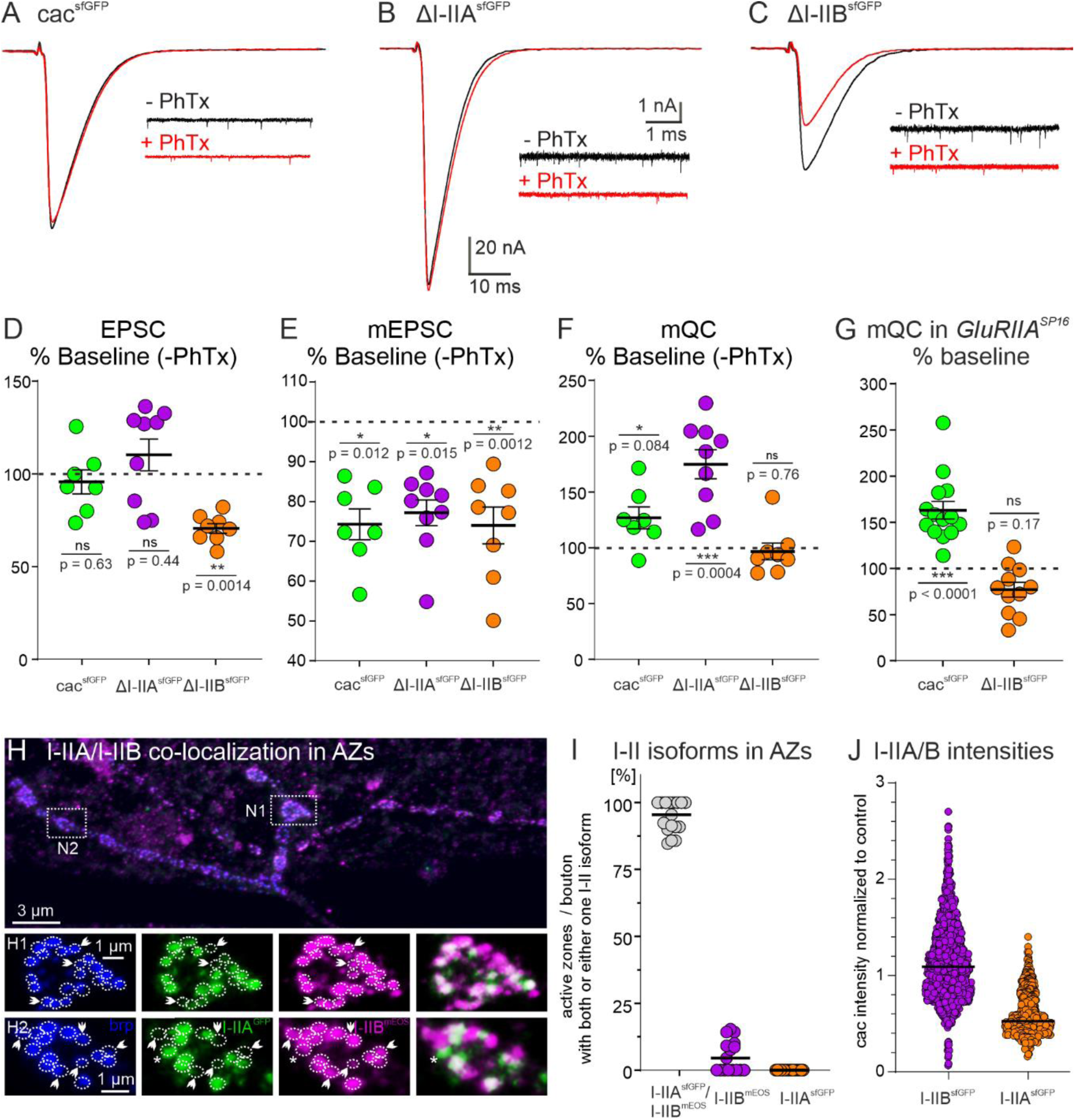
Removal of I-IIB impairs presynaptic homeostatic potentiation. **(A-C)** Acute presynaptic homeostatic potentiation (PHP) can be induced in control (cac^sfGFP^) and *ΔI-IIA* but not in *ΔI-IIB* animals by bath application of the glutamate IIA receptor (GluRIIA) blocker philanthotoxin (PhTx) (A-C; black traces: without PhTx, red traces after PhTx application). **(D-F)** Quantification of EPSC amplitude, mEPSC amplitude, and mean quantal content (mQC) is shown as % change in PhTx treated animals to untreated control within genotypes (% Baseline (−PhTx). Untreated controls are set to 100 % (dotted line in **D-F**). EPSC amplitudes in cac^sfGFP^ and *ΔI-IIA* animals are not significantly affected (**D**, cac^sfGFP^, green p=0.63, *ΔI-IIA*, purple p=0.44), while the EPSC amplitude in *ΔI-IIB* animals is significantly reduced after PhTx treatment (**D**, orange, p=0.0014). However, reduction of mEPSC amplitude in all genotypes shows successful block of GluRIIA compared to the respective control (**E**, % change to untreated control within genotype; cac^sfGFP^, green p=0.012; *ΔI-IIA*, purple p=0.0015, *ΔI-IIB*, orange p=0.0012). Accordingly, mean quantal content (mQC) is increased in cac^sfGFP^ and *ΔI-IIA* animals but not in *ΔI-IIB* animals (**F**, cac^sfGFP^, green p=0.084; *ΔI-IIA*, purple p=0.0004, *ΔI-IIB* p=0.76). **(G)** PHP maintenance is typically assessed in *GluRIIA* mutants. PHP maintenance is observed in cac^sfGFP^ animals because mQC is increased in a *GluRIIA^SP16^* mutant background (*GluRIIA^SP16^*; green, p<0.0001). By contrast, in *ΔI-IIB* animals the *GluRIIA* mutant background does not cause an increase in mQC (orange p=0.17). All pairwise comparisons in **(D-G)** use unpaired Student’s T-test between untreated and treated condition within each genotype. **(H)** In animals that are transheterozygous for the removal of *I-IIA* and the removal of *I-IIB*, and carry GFP-tagged I-IIA and mEOS4b-tagged I-IIB cacophony channels (I-IIA^sfGFP^/I-IIB^mEOS^), triple immunolabel for the AZ marker brp, GFP, and mEOS4b show that most AZs (blue) but not all (white arrow heads in H1, H2 and asterisk in H2) contain both, GFP-tagged I-IIA (green) and mEOS4b-tagged I-IIB (magenta) channels. Magenta label in the overlay (H1, H2, right column, arrow heads) indicates AZs with only I-IIB channels, green label in the overlay (H2, right column, asterisk) indicates one AZ with only I-IIA channels. **(I)** Quantification shows that >95 % of all brp positive AZs contain I-IIA and I-IIB channels (gray dots), few AZs (∼5 %) contain only I-IIB (= *ΔI-IIA*; purple), and almost no AZ contains only I-IIA (= *ΔI-IIB*; orange). **(J)** Relative fluorescence intensity of sfGFP-tagged I-IIB channels (= *ΔI-IIA*; purple) and sfGFP-tagged I-IIA channels (= *ΔI-IIB*; orange). For assessment of I-IIB channels, tagged channels were expressed transheterozygously over untagged I-IIA channels (*ΔI-IIA^sfGFP^/ΔI-IIB^no^ ^tag^*; purple) and *vice versa* (ΔI-*IIB^sfGFP^/ΔI-IIA^no^ ^tag^*; orange). Quantification of relative fluorescence intensity compared to transheterozygous control (*cac^sfGFP^/cac^no^ ^tag^*) reveals an expression ratio of I-IIB to I-IIA channels of 2:1.

Given that synapses that contain only cac isoforms without the *I-IIB* exon show a lower release probability, more variable paired pulse ratios and synaptic depression, and an inability for compensatory increases in mean quantal content, the question arises whether normal synapses contain the cac^I-IIA^ channels at all. This can be tested in transheterozygous *ΔI-IIA^GFP^/ΔI-IIB^mEOS4b^* animals, which show normal EPSC amplitudes as well as mEPSC amplitudes and frequencies (Figs. 4J-L). Co-labeling of AZs with anti-brp (Fig. 7H, blue) in animals that contain cac channels without I-IIB but with GFP-tagged I-IIA (Fig. 7H, green) on one chromosome and cac channels without I-IIA but with mEOS4b-tagged I-IIB (Fig. 7H, magenta) indicates that all AZs contain cac with I-IIB, some AZs contain only I-IIB channel (Fig. 7H1, H2, white arrow heads), most contain both I-IIB and I-IIA, but very few AZs contain cac channels with only I-IIA (Fig. 7H2, asterisk). Quantification reveals that ∼95 % of all AZs contain cac channels with both, the I-IIA and the I-IIB exon, less than 5 % of the AZs contain only cac with the I-IIB exon and almost no AZ contains only cac with the I-IIA exon (Fig. 7I). These data show that the vast majority of presynaptic AZs house a mixture of I-IIA and I-IIB cac. To test whether there is a consistent average ratio of I-IIA and I-IIB channels in presynaptic AZs we measured the cac puncta fluorescence intensities for heterozygous cac^sfGFP^/cac, cac^I-IIA^ ^sfGFP^/cac^I-IIB^, and cac^I-IIB^ ^sfGFP^/cacI-IIA animals. This way intensity is always measured from cac puncta with the same GFP tag. Normalizing all values to the intensities obtained in AZs from heterozygous cac^sfGFP^/cac controls reveals a consistent ratio 2:1 in the relative intensities of I-IIB and I-IIA across junctions and animals (Fig. 7J). This is consistent with the counts in our spt-PALM analysis (see Fig. 5F) and indicates on average roughly twice as many I-IIB as compared to I-IIA channels across AZs.

## Discussion

The Drosophila *Ca*_v_*2* homolog is named *cacophony* and contains two mutually exclusive splice sites that do not exist in vertebrate *Ca*_v_*2* VGCC genes. Our data show that alternative splicing at these sites substantially increases Drosophila cac functional heterogeneity. We report that the first mutually exclusive exon, that is located in the fourth transmembrane domain of the first homologous repeat (*IS4*), affects cac biophysical properties and is decisive as to whether the channels localize to presynaptic active zones (AZs) and thus participate in fast synaptic transmission. By contrast, mutually exclusive splicing at the second site encoding the intracellular linker between the first and the second homologous repeat (I-II) does not affect cac presynaptic AZ localization, but instead, fine-tunes multiple different aspects of presynaptic function. In vertebrates, substantial functional synaptic heterogeneity can result from different combinations of Ca_v_2.1, Ca_v_2.2, and/or Ca_v_2.3 in the presynaptic AZ (reviewed in Zhang et al. 2022). In Drosophila, the collective functions of mammalian Ca_v_2.1, Ca_v_2.2, and Ca_v_2.3 must be portrayed by one *Ca*_v_*2* gene. Mutually exclusive splicing at the *IS4* and *I-II* sites might have been a different evolutionary strategy to create cac mediated functional synaptic heterogeneity. However, alternative splicing increases functional diversity also in mammalian Ca_v_2 channels. Although the mutually exclusive splice site in the S4 segment of the first homologous repeat (IS4) is not present in vertebrate Ca_v_ channels, alternative splicing in the extracellular linker region between S3 and S4 is at a position to potentially change voltage sensor properties (Bezanilla 2002). Alternative splice sites in rat Ca_v_2.1 exon 24 (homologous repeat III) and in exon 31 (homologous repeat IV) within the S3-S4 loop modulate channel pharmacology, such as differences in the sensitivity of Ca_v_2.1 to Agatoxin. Alternative splicing is thus a potential cause for the different pharmacological profiles of P– and Q-channels (both Ca_v_2.1; Bourinet et al. 1999). Moreover, the intracellular loop connecting homologous repeats I and II is encoded by 3-5 exons and provides strong interaction with G_βγ_-subunits (Herlitze et al. 1997). In Ca_v_2.1 channels, binding to G_βγ_ –subunits is potentially modulated by alternative splicing of exon 10 (Bourinet et al. 1999). Moreover, whole cell currents of splice forms α1A-a (no Valine at position 421) and α1A-b (with Valine) represent alternative variants for the I-II intracellular loop in rat Ca_v_2.1 and Ca_v_2.2 channels. While α1A-a exhibits fast inactivation and more negative activation, α1A-b has delayed inactivation and a positive shift in the IV-curve (Bourinet et al. 1999). This is phenotypically similar to what we find for the mutually exclusive exons at the IS4 site, in which IS4B mediates high voltage activated cacophony currents while IS4A channels activate at more negative potentials and show transient current (Fig. 3; see also Ryglewski et al. 2012). Furthermore, altered Ca_β_ interaction have been shown for splice isoforms in loop I-II (Bourinet et al. 1999), similar to what we suspect for the I-II site in cacophony. Finally, in mammalian VGCCs, the C-terminus presents a large splicing hub affecting channel function as well as coupling distance to other proteins. Taken together, Ca_v_2 channel diversity is greatly enhanced by alternative splicing also in vertebrates, but the specific two mutually exclusive exon pairs investigated here are not present in vertebrate *Ca*_v_*2* genes. Below, we will discuss the consequences of Drosophila *cac* splicing at these sites for presynaptic function.

### Exon IS4B is required for sustained HVA current and presynaptic AZ localization

Mutually exclusive splicing in the 4^th^ transmembrane domain of the first homologous repeat (*IS4*) yields either *IS4A* or *IS4B*. Our data show that the *IS4B* exon is required for presynaptic AZ localization of cac and for evoked synaptic transmission at a fast glutamatergic synapse, the Drosophila NMJ. Accordingly, excision of the *IS4B* exon is embryonic lethal. This is in accordance with the finding that the only cac knock-in construct that has ever been reported to rescue lethality of Drosophila *cac* null mutants contains the *IS4B* exon (*UAS-cac1*; Kawasaki et al. 2004). The other alternative exon at the *IS4* site, *IS4A*, is not sufficient to mediate presynaptic AZ localization, does not contribute to evoked synaptic transmission at fast synapses, but it gives rise to functional cacophony channels that localize to other neuron types and neuronal compartments. Therefore, the absence of cac^IS4A^ isoforms from presynaptic AZs at the NMJ is not a consequence of general channel degradation upon excision of *IS4B*, but of exon specific channel localization. In fact, cac^IS4A^ isoforms are expressed sparsely but in stereotypical patterns in the larval brain and VNC (see suppl. Fig. 1). Moreover, in the absence of IS4B isoforms, cac^IS4A^ channels mediate somatodendritic calcium current in adult motoneurons. By contrast, cac^IS4A^ isoforms are not detectable in presynaptic boutons at the larval NMJ. Accordingly, animals with excision of *IS4A* show normal neuromuscular transmission.

Our voltage clamp recordings from motoneuron somata show that splicing in the 4^th^ transmembrane domain, where the voltage sensor is located, affects cac activation voltage. Removing the *IS4B* exon virtually abolishes fast activating, sustained cac mediated HVA current. We infer that *IS4B* containing cac mediate fast activating, sustained HVA current also in presynaptic AZs, although we cannot exclude the possibility that cac^IS4B^ interacts with different accessory calcium channel subunits or with different other proteins to give rise to macroscopically different calcium currents, depending on whether the cac α_1_-subunit localizes to presynaptic AZs, or to other subcellular compartments. However, fast activating, sustained HVA current is in accordance with the demands of the Drosophila larval NMJ that transmits burst of ∼200 ms duration with action potential frequencies of ∼120 Hz during crawling (Kadas et al. 2017). Fast voltage gated activation of sustained HVA calcium current with fast inactivation upon repolarization is useful at large intraburst firing frequencies without excessive cac inactivation. Our somatic voltage clamp recordings further demonstrate that *IS4B* is not only essential for evoked synaptic transmission, but cac with the *IS4B* exon can also give rise to somatodendritic HVA current. Therefore, the *IS4B* exon is essential for presynaptic function but it can also give rise to cac currents in other neuronal compartments and it is abundantly expressed throughout VNC neuropils. By contrast, the cac^IS4A^ channels are not expressed in NMJ presynaptic terminals, but they show a reproducible sparse expression pattern in some VNC regions, thus likely mediating some distinctly different functions that are yet uncharacterized. An abundant function of IS4B channels in presynaptic AZs of fast chemical synapses is in line with strong protein detection in Westerns from brains, whereas the sparse expression of IS4A in distinct sub-regions of the CNS is in line with a faint band in Western Blots.

### I-II exon alternative splicing fine-tunes presynaptic function

In contrast to the *IS4* exon, mutually exclusive splicing in the intracellular loop between the first and the second homologous repeats (I-II) does not affect cac localization in AZs at the NMJ. In fact, >95 % of all presynaptic AZs contain both, cac isoforms with I-IIA and cac isoforms with I-IIB. Single on-locus alternative exon removal at the *I-II* site allows the study of presynaptic cac channel function with only one of the mutually exclusive I-II exons. Removal of *I-IIA* (*ΔI-IIA*) leaves 8 isoforms that contain *I-IIB*. This does not affect cac localization or channel number in presynaptic AZs as compared to control. However, half amplitude width of evoked EPSCs is significantly decreased whereas median amplitude of evoked EPSCs is increased from ∼115 nA to ∼130 nA, although the amplitude increase is statistically just not significant (one sided Mann Whitney U-test, p = 0.078). Together a slight amplitude increase and a decreased half width likely cancel out changes in EPSC charge. Similar channel numbers and localizations but altered EPSC amplitudes and shapes could potentially be caused by different cac properties in the absence of *I-IIA* as compared to control. Specifically, decreased EPSC half width could be caused by significantly faster channel inactivation kinetics, and slightly increased single channel conductance may increase EPSC amplitude. Alternatively, altered EPSC width and amplitude could also be caused by changes in presynaptic action potential shape or postsynaptic glutamate receptor properties or compositions. Although we found no difference in GluRIIA abundance between *ΔI-IIA* and control, we cannot exclude other changes in postsynaptic receptor fields. However, given that *I-IIA* excision primarily affects the relative abundance of I-IIA and I-IIB cac in the presynaptic AZ but not mean channel number, given that postsynaptic VGCCs in muscle are encoded by the Drosophila *Ca*_v_*1* and *Ca*_v_*3* homologs, and given that cac channels localize to axon terminal AZs, we consider altered properties of different cac isoforms a possible explanation for altered EPSC properties upon *I-IIA* excision. During repetitive firing, the median increase of EPSC amplitude by ∼10 % is potentially counteracted by the significant decrease in EPSC half amplitude width by ∼25 %, so that neither paired pulse ratios, nor synaptic depression show significant differences between control and *ΔI-IIA*. Even for stimulus trains that mimic intraburst motoneuron activity as observed during restrained crawling in semi-intact preparations (Kadas et al. 2017), there is no difference in charge transfer from the motoneuron axon terminal to the postsynaptic muscle cell between *ΔI-IIA* and control. Surprisingly, crawling is significantly affected by the removal of *I-IIA*, in that the animals show a significantly increased mean crawling speed but no significant change in the number of stops. Given that the presynaptic function at the NMJ is not strongly altered upon *I-IIA* excision, and that *I-IIA* likely mediates also cac functions outside presynaptic AZs (see above) and in other neuron types than motoneurons, and that the muscle calcium current is mediated by *Ca*_v_*1* and *Ca*_v_*3*, the effects of *I-IIA* excision of increasing crawling speed is unlikely caused by altered pre-or postsynaptic function at the NMJ. We judge it more likely that excision of *I-IIA* has multiple effects on sensory and pre-motor processing, but identification of these functions is beyond the scope of this study.

Removal of *I-IIB* (*ΔI-IIB*) leaves 10 cac isoforms that contain I-IIA. This does not affect presynaptic AZ localization of cac or the proximity to the AZ scaffold protein brp (Kittel et al. 2006), but it significantly reduces P_r_ by ∼ 50 %. This bisection in P_r_ is correlated with a 50 % reduction in the number of cac channels in AZs from about 10 to about 5, as determined by tagged cac fluorescence intensity measurement in fixed specimen and by live sptPALM counting of cac^mEOS4b^ (Heck et al. 2019; Ghelani et al. 2023). These data suggest that P_r_ is nearly linearly related to the number of VGCCs in the presynaptic AZ, at least at 0.5 mM external calcium which we mainly used for this study. Please note that the relation between external calcium and P_r_ at the Drosophila NMJ has been reported non-linear at external calcium concentrations above 1.5 mM, but linear between 0.5 and 1.5 mM (Weyhersmüller et al. 2011). Accordingly, at 1.5 mM external calcium a linear correlation between P_r_ and tagged cac channel fluorescence intensity has recently been reported (Medeiros et al. 2024). This also corresponds to observations at the mammalian calyx of Held, where the number of AZ VGCCs correlates linearly with P_r_, and moreover, influences whether subsequent release is depressed or facilitated (Sheng et al. 2012). Similarly, in synapses with fewer cac channels upon *I-IIB* excision, we find a significant decrease in P_r_ along with a significant reduction in paired pulse ratio (PPR). The effects on PPR can by fully attributed to reduced channel numbers in presynaptic AZs upon *I-IIB* excision, because they can be fully rescued by triturating the external calcium concentration so that the first pulse amplitude matches that of controls with normal P_r_. Therefore, regulation of VGCC number in presynaptic AZs may be a conserved mechanism to tune P_r_ and short-term plasticity from flies to mammals.

At the Drosophila NMJ a steep gradient of synaptic transmission amplitude exists along the motoneuron axons over their target muscle fibers, with the highest presynaptic Ca^2+^ influx in most distal presynaptic sites along the axon. This has been interpreted as gradient control of P_r_ along the axon of the same neuron (Guerrero et al. 2005), which is at least in part regulated by the balance of different release enhancing and suppressing proteins at proximal versus distal release sites, such as complexin (Newman et al. 2022). Another potential means to regulate P_r_ at different release sites of the same neuron could be uneven ratios of I-IIA and I-IIB cac isoforms, which requires additional analysis. The prediction would be that the more distal the release site the more I-IIB channels are expressed. Given that these have a highly conserved Ca_β_ binding motif, targeting to distal release sites might be promoted in comparison to I-IIA channels with a less conserved Ca_β_ binding motif (Smith et al. 1998).

The reduction in AZ cac number upon removal of *I-IIB* has three additional important functional consequences. First, reduced P_r_ and charge transfer across the NMJ during crawling-like motoneuron bursting patterns are reflected in a significant decrease in mean crawling speed and a highly significant increase in the number of stops, whereas maximum crawling speed is not affected. Unaffected maximum speed at significantly reduced P_r_ indicates that the charge transfer at high motoneuron intrabust firing frequencies is above the one needed for maximum muscle contraction. Moreover, mean speed is significantly but just slightly decreased upon removal of *I-IIB* and a highly significant reduction in P_r_, whereas the number of stops is increased highly significantly by ∼50 %. It seems likely that large effects on the continuation of motor behavior but only mild effects on the speed are caused not only by effects on synaptic transmission at the NMJ but also by effects on other neuronal compartments or on other neurons in sensory or premotor circuitry. This interpretation would be in accordance with the finding that cac currents are also measured from the somatodendritic domain of pupal and adult Drosophila (this study) and that larval Drosophila motoneuron excitability as measured by I/F relationships are altered upon *cac-*RNAi (Worrell, Levine, 2008). Second, removal of *I-IIB* increases the variability of paired pulse ratio and the time course of synaptic depression, but this is solely due to reduced channel numbers upon *I-II* excision, because the effect is rescued by increasing the external calcium concentration. It seems plausible that mechanism with probabilistic features, such as cac activation/inactivation/de-inactivation as well as Ca_v_2 channel mobility in AZs (Heck et al. 2019; Ghelani et al. 2023) exert a higher impact with fewer channels. Therefore, increasing variability of presynaptic function might be another consequence of reduced AZ cac number upon *I-IIB* excision. And third, removal of *I-IIB* abolished the ability of both initiation and maintenance of presynaptic homeostatic potentiation (PHP), which in turn, requires an upregulation of the number of cac and of brp molecules in the presynaptic AZ (Ghelani et al. 2023; Gratz et al. 2019). Similarly, PHP is also blocked in cac hypomorphic mutants which also reduce EPSC amplitude, likely due to reduced calcium influx (Frank et al., 2006), thus indicating that fast PHP induction might not be possible with reduced calcium conductance in AZs. Similarly, increased calcium influx into the presynaptic AZ after induction of PHP (Müller, Davis 2012; 2015) fits with an increase in cac channel number. Increasing the number of cac^I-IIA^ in AZs as a compensatory response to reduced postsynaptic receptor function seems not likely with reduced Ca_β_ binding affinity, or in the face of fewer channels to start with, or for additional reasons, which will require additional studies.

In summary, our study suggests mutually exclusive splicing at two *cac* splice sites, that do not exist in mammals, as an alternative strategy to increase functional heterogeneity at the presynaptic AZ of fast chemical synapses, so that the Drosophila *Ca*_v_*2* homolog *cacophony* may portrait some of the functional heterogeneity that arises in mammals from the combinatorial usage of Ca_v_2.1, Ca_v_2.2, and Ca_v_2.3. Splicing at the first Drosophila mutually exclusive site directs cac to different subcellular compartments and tunes cac biophysical properties. Splicing at the second mutually exclusive site does not direct cac to different subcellular compartments, but it fine tunes multiple aspects of presynaptic function by changing presynaptic AZ channel numbers or ratios between cac splice variants.

## Methods

### Generation of CRISPR flies

Exon excision was performed via the CRISPR/Cas9 method (Doudna and Charpentier, 2014; Sternberg and Doudna, 2015). Cacophony is located on the X-chromosome in Drosophila. Cac^sfGFP^ exon out flies were generated by crossing female virgin flies expressing super folder GFP (sfGFP)-tagged cac (C cac^sfGFP^, Gratz et al. 2019) channels along with the Cas9 enzyme under the control of the germ line active *nanos*-promoter (*nos-cas9*, Bloomington stock center #78781) to male flies expressing a *gRNA* transgene under the control of the germ line active *U6*-promoter. The *gRNA* sequences were designed such to specifically target sequences flanking the exon to be excised (see table 1). CRISPR events take place as soon as both *nos-cas9* as well as *U6-gRNA* transgenes are present in the same fly, no matter whether these flies are male or female. For simplicity, for excision of exons *IS4A*, *I-IIA*, and *I-IIB*, male progeny were collected as such flies are hemizygous vital, whereas for excision of *IS4B*, females were collected because removal of *IS4B* is lethal. Flies were then back-crossed into suitable balancer strains to keep the X-chromosome with the putative cac exon excision and to be able to follow out-crossing of *nos-cas9* or *U6-gRNA* transgenes. Successful exon excision was confirmed with single fly genomic PCR with suitable primers and Taq DNA polymerase (New England Biolabs, #M0267S) (for cycler (Biorad T100 thermo cycler) settings and primers see tables 2 and 3) and subsequent 1% agarose gel electrophoresis. Gene sequence around the excision was confirmed with next generation sequencing (StarSeq, University of Mainz Campus with the exon out verification primers, see table 3). Gels were run at 100 V for 45 minutes. Primers were obtained from Integrated DNA Technologies, Germany. To minimize contamination of exon out fly stocks, successful excision mutants were cantonized for at least 5 generations to enhance the likelihood of outcrossing of undesired mutations due to CRISPR off-target events. To clean up the X-chromosome itself on which the desired CRISPR event took place, flies were subjected to recombination with Canton S flies. Lack of the desired exon was then re-confirmed by PCR again.

**Table 1:**
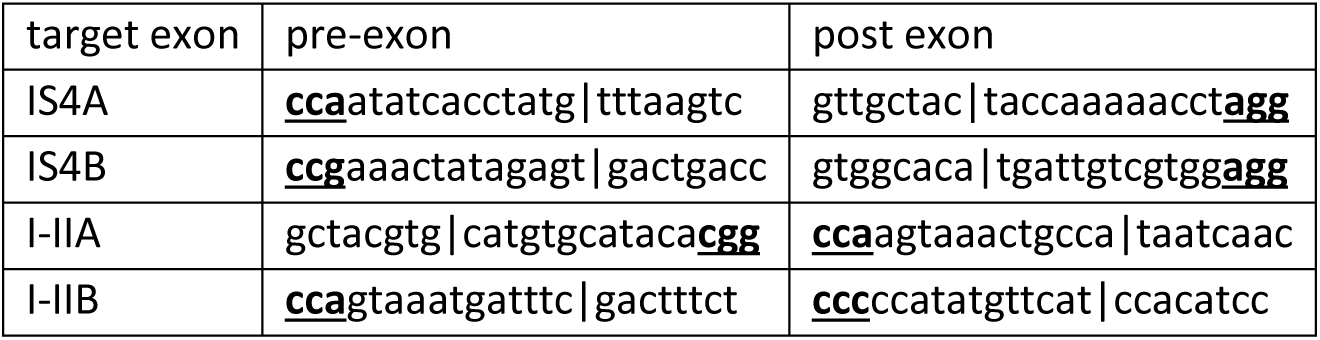
gRNA sequences used for cas9 target site. Vertical lines depict intended break points. Bold and underlined nucleotides indicated PAM (protospacer adjacent motif) sequences.

**Table 2:** Cycler settings for exon out verification.

| Step | temperature |  | duration |
| --- | --- | --- | --- |
| 1 | 95°C |  | 3:00 min |
| 2 | 95°C |  | 0:30 min |
| 3 | 57°C | -1°C per cycle | 0:45 min |
| 4 | 68°C |  | 1:30 min |
| 5 | Go to step 2 | 10x |  |
| 6 | 95°C |  | 0:30 min |
| 7 | 52°C |  | 0:45 min |
| 8 | 68°C |  | 1:30 min |
| 9 | Go to step 6 | 24x |  |
| 10 | 68°C |  | 5:00 min |
| 11 | 4°C |  | ∞ |

**Table 3:**
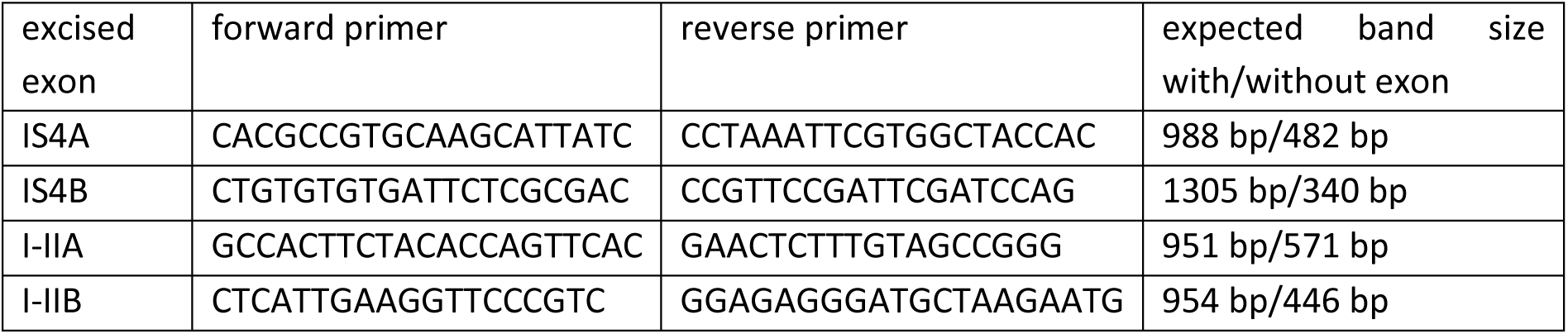
Exon out verification primers.

### Generation of gRNA transgenic flies

First, the eligibility for Cas9 cleavage was assessed, core properties are the protospacer adjacent motif (PAM) NGG flanking the 20 bp *gRNA* (see table 1) to facilitate site recognition, and a 5’ G as the vector used for *gRNA* expression later used the *U6*-promoter. Moreover, CRISPR sites must not disrupt recognition sites for the splicing machinery. For this, online tools were used (CRISPR target finder: Gratz et al. 2014, http://targetfinder.flycrispr.neuro.brown.edu/index.php and Cas9 target finder: https://shigen.nig.ac.jp/fly/nigfly/cas9/cas9TargetFinder.jsp). Sites for double strand breaks were picked at a distance from the intron/exon borders to not affect splice acceptor sites. Furthermore, sites known to impact splice efficiency (Blanchette et al. 2005; Brooks et al. 2011) were avoided. In addition, sites for minimal loss of total genomic region were determined and the *gRNA* sequences with as few as possible predicted off-target cleavage sites were selected.

### Fly rearing

Flies were kept at 25°C in 25 mm diameter plastic vials with mite proof foam stoppers on a cornmeal, yeast, agar, glucose diet on a 12 hr light/dark regimen. Wandering L3 larvae were collected directly from food vials. 2-day old adult flies were collected from their food vials and placed on ice in pre-chilled empty food vials for no more than 1 minute prior to dissection (for patch clamp recordings).

### Flies

For electrophysiological recordings as well as for immunohistochemical labeling, sfGFP-tagged cac flies were used. First, validity of sfGFP-tagged cac channel flies was confirmed by comparing w^+^ TI{TI}cac^sfGFP-N^ flies with the wildtype strain Canton special (Canton S). After that, the control strain for all sfGFP-tagged exon out flies was w^+^ TI{TI}cac^sfGFP-N^

Exon out flies were w^+^ TI{TI}cac^sfGFP-N^ ^ΔIS4A^, w^+^ TI{TI}cac^sfGFP-N^ ^ΔIS4B^/FM7c P{2x Tb-RFP}, w^+^ TI{TI}cac^sfGFP-N^ ^ΔI-IIA^, w^+^ TI{TI}cac^sfGFP-N^ ^ΔI-IIB^. All CRISPR fly strains were originally white mutant. To reduce the likelihood of accumulation of off-target effects resulting from CRISPR events, we replaced the mutant white gene on the X-chromosome (on which *cacophony* is also located) by a Canton S derived wildtype white gene from our Canton S lab stock that we also used as control (see above). In addition, after CRISPR events, flies were crossed to remove nos-Cas9 and gRNA transgenes on the 2^nd^ chromosome and replaced these chromosomes by ones without transgenes or other known mutations. This first removed transgenes that were needed for CRISPR induction and second, this also removed possible off target events on the 2^nd^ chromosome. In the process, 3^rd^ and 4^th^ chromosomes were replaced automatically as well. We back crossed with Canton S strains for at least 5 generations. Thus, off-target chromosomal aberrations were minimized by recombination on the X-chromosome and by replacement of all other chromosomes by out-crossing.

For channel counting and for assessment of cac^I-IIA^ and cac^I-IIB^ expression in AZs (both see below), we used exon out flies that carried mEOS4b endogenously directly after the start codon of *cacophony* (Ghelani et al. 2023). mEOS4b-tagged exon out fly strains were: w* TI{TI}cac^mEOS4b-N^ w* TI{TI}cac^mEOS4b-N^ ^ΔI/IIA^, w* TI{TI}cac^mEOS4b-N^ ^ΔI/IIB^. All mEOS4b-tagged fly strains were white mutant.

For GluRIIA immunohistochemistry, transgenic flies expressing GluRIIA^GFP^ under a native promoter were used (Rasse et al. 2005).

### Western Blots

For proof of protein expression after exon excision, Western Blots were performed with detection of the N-terminal sfGFP tag (due to lack of cacophony antibodies). As loading control, β-actin was used. Cacophony bands were expected above 250 kDa (cacophony plus GFP) and actin was expected at 42 kDa. For viable exon excision mutants (cac^sfGFP^ as control, cac^sfGFP^ ^ΔIS4A^, cac^sfGFP^ ^ΔI-IIA^, cac^sfGFP^ ^ΔI-IIB^), 10 adult male brains per lane were prepared, for lethal exon excisions (Canton S as control, cac^sfGFP^/+, cac^sfGFP^ ^ΔIS4B^/+), 20 adult female brains of heterozygous animals were used. Brains were dissected with forceps one by one and collected in 23 µl squishing buffer (composition below) in 0.5 ml low bind sample tubes on ice. When 10 or 20 brains, respectively, were collected, samples were squished with a sterile pestle using a motorized squishing device. After squishing, 3 µl sample buffer (composition below) were added to the homogenized samples, which were subsequently boiled at 95°C for 5 min. Samples were then stored at –30°C until use. For loading, samples were taken out of the freezer, boiled for 3 minutes at 95°C and directly loaded into a 10 well Mini-Protean TGX Precast Gel with a 4-15% polyacrylamide gel gradient with 50 µl volume per well (Biorad, Cat#456-1084). The gel was run in running buffer (composition below) for 2h 40 minutes at 80 V. As protein marker, 8 µl Spectra Multicolor High Range Ladder (Thermo Scientific, Cat# 26625) was used. Transfer was done in transfer buffer (composition below) on nitrocellulose membrane overnight at 35 V with a cool block and the setup sitting in an ice box. Next day, the nitrocellulose membrane was washed with TBS-Tween20 and then subjected to blocking with Intercept T20 Antibody Diluent (LI-COR, Cat#927-65001) diluted 1:1 with 0.1M PBS for 2 hrs. Then the nitrocellulose membrane was cut horizontally to subject the pieces with the expected cacophony bands and the expected loading control (β-actin) separately to antibody labeling. This was followed by primary antibody incubation with 1:500 polyclonal rabbit anti-GFP antibody (Thermo Fisher Scientific, Cat# A11122) or 1:10,000 monoclonal mouse anti-actin (DSHB, JLA20) diluted in Intercept T20 Antibody Diluent (see above), first for 2 hrs at room temperature and then for two nights at 4°C. Then antibody solution was removed and membrane washed 3× 15 minutes with TBS-Tween20. This was followed by incubation with secondary antibodies diluted in Intercept T20 Antibody Diluent for 2 hrs at room temperature in the dark with IRDye 680 donkey anti-rabbit (LI-COR, Cat# 926-68073) at 1:10,000 to detect GFP-tagged cacophony bands, or IRDye 800 donkey anti-mouse 1:10,000 to detect β-actin, respectively. This was followed by 2× 15 minutes washes with TBS-Tween20 and then 1× 15 minutes with 0.1M PBS to reduce background. Bands were detected with a LI-COR Odyssey Fc Imaging System with 30 second or 2 minutes exposure times.

### Western Blot solutions

Lyse buffer: 25 ml 4xTris-HCl/SDS pH 6.8, 20 ml glycerol, 4 g SDS, 1 mg bromophenol blue.

6x sample buffer (10 ml): 0.3 M Tris-HCl pH 6.8, 8% SDS, 50% glycerol, 0.25% bromophenol blue. 8% mercapto-ethanol was added fresh directly before use.

10x SDS running buffer: 30 g Tris Base (pH 8.3), 150 g glycine, 10 g SDS. Add ddH_2_O to 1000 ml total volume.

Transfer buffer: 15.5 g Tris Base, 72 g glycine, 1000 ml methanol, 5000 ml ddH_2_O.

### Dissection for electrophysiology and immunohistochemical labeling

Third instar larvae were dissected in HL3.1 saline (composition below). Larvae were fixed dorsal side up to a Sylgard (Sylgard 184, DowCorning) coated lid of a 35 mm Falcon dish with two insect minuten pins through the mouth hook and through the tail. After covering the larva with saline (composition below), the body wall was cut along the dorsal midline. To obtain a filet preparation, the body wall was spread laterally and fixed with two minuten pins on each side. Gut and trachea were removed as well as the ventral nerve cord. For electrophysiology from the larval NMJ (see below: electrophysiology), the motor nerves were left long, for immunohistochemical labeling (see below: immunohistochemistry), the motor nerves were cut very short.

Adult flies were dissected in normal saline (composition below – but calcium current recordings were performed with different external solution – composition and electrophysiology see below). Adult two-day old female flies were anaesthetized briefly on ice in pre-chilled empty fly rearing vials. Then legs and wings were removed and the fly fixed dorsal side up in a Sylgard coated lid of a 35 mm Falcon dish with two insect minute pins through the head and through the tip of the abdomen. After covering the fly with saline (composition below), the cuticle was cut along the dorsal midline. To expose the ventral nerve cord with the motoneurons, the thoracic cuticle with the cut wing depressor muscle (DLM) was spread laterally and pinned down with one minuten pin on either side. After removal of gut, esophagus, and salivary glands, the ventral nerve cord was exposed.

### Electrophysiology

All electrophysiological recordings were carried out at room temperature (∼ 22°C). For TEVC recording microelectrodes were pulled with a Sutter Flaming Brown P97 microelectrode puller from borosilicate glass capillaries with filament, with inner diameter of 0.5 mm, and an outer diameter of 1 mm (Sutter BF100-50-10 or World Precision Instruments 1B100F-4). Patch pipettes were pulled with a Narishige PC10 electrode puller from borosilicate glass capillaries without filament, with an inner diameter of 1 mm, and an outer diameter of 1.5 mm (World Precision Instruments PG52151-4).

### TEVC of muscle 6 or muscle 12 in the larval body wall

For two electrode voltage clamp (TEVC) recordings, dissected third instar larvae (see dissection) were placed on an upright Olympus BX51WI fixed stage microscope with a 20x water dipping lens (Zeiss LD A-Plan). The motor nerves of either segment A2 or A3 on the left or the right side was sucked into a glass microelectrode filled with saline with an individually broken tip. After placing the recording electrodes filled with 3 M potassium chloride (KCl) and the ground wire (chlorinated silver) into the recording solution (HL3.1 saline, composition see below), both electrodes were nulled against the ground electrode (bridge mode) using an Axoclamp 2B intracellular amplifier (Molecular Devices). Then the muscle (M6 or M12) cell was first impaled with the larger tip current passing glass microelectrode (∼8 – 13 MΩ) directly followed at a small distance by the smaller tip recording electrode (∼20 – 30 MΩ). If both electrodes recorded a membrane potential of at least –50 mV without differing by more than 3 mV, RMP balance was adjusted and then TEVC established by pressing the respective knob on the amplifier. Gain was set between 8 and 25. Anti-alias filter was set to ½ the sampling rate to minimize noise. Then the desired command potential was set, mostly at –70 mV. Holding current needed to keep the membrane potential at the desired –70 mV had to be below 4 nA, otherwise recordings were discarded. Data were digitized at 50 kHz (Digidata 1550B, Molecular Devices) and recorded with pClamp 11.1.0.23 software package (Molecular Devices). Data were filtered offline with a lowpass Gaussian Filter with a –3 dB cutoff frequency of 360 Hz.

The suction electrode with the motor nerve was connected to a pulse generator (A-M Systems Model 2100) to deliver low voltage (up to 10 V) 0.1 ms duration stimulations at varying patterns. Stimulation voltage was adjusted slowly to be sure that both motor axons innervating the recorded muscles were stimulated. This was determined unequivocally by the transmission amplitude, which is distinctly smaller if just one motor axon is stimulated as compared to stimulation of both motor axons.

For recordings of miniature excitatory postsynaptic currents (mEPSCs), spontaneous occurrence of mEPSCs was recorded for one minute without stimulation. For single evoked excitatory postsynaptic currents (EPSCs), the motor nerve was stimulated at 0.1 Hz frequency for 50 seconds and the amplitude was recorded. Mean quantal content was determined by dividing the mean amplitude of several EPSCs of one recording by the mean amplitude of several mEPSC of the same recording. For short term plasticity, paired pulses (PPs) were recorded by stimulation of the motor nerve twice with varying interspike intervals (IPIs) of 10 ms, 15 ms, 20 ms, 25 ms, 30 ms, 50 ms, and 100 ms (applied in that order with intersweep intervals of 10 s). Paired pulse ratios (PPRs) were calculated by dividing the amplitude of the second pulse by that of the first pulse. Mean PPR values are based on the PPRs of each sweep and then averaged and also by first averaging the amplitudes of all first and all second pulses separately, to then divide the mean of the second by the mean of the first amplitude. In contrast to work on pyramidal neurons (Kim and Alger 2001) in our data PPRs not different depending on these two analysis schemes. Short term facilitation or short term depression were induced by 60 s stimulation at 1 Hz or by 1s stimulation at 10 Hz or by 200 ms stimulation at 60 Hz and 120 Hz stimulation frequency. For each animal the amplitudes of all subsequent EPSCs in each train were plotted over time and fitted with a single exponential. For depression at 1 and 10 Hz, we used one train per animal, and 5-6 animals per genotype. Motoneuron burst firing as occurring during crawling (Kadas et al. 2017) was simulated by higher frequency stimulation for 200 ms at 60 Hz and 120 Hz.

Presynaptic homeostatic potentiation (PHP) was induced either acutely by application of the bee wolf toxin Philanthotoxin 433 or by using animals carrying a null mutation in the GluRIIA glutamate receptor that is expressed in the muscle membrane. PhTx (10 µM) is applied in calcium-free HL3.1 saline (composition see below) after dissection but before mounting the preparation on the microscope stage. For the toxin to work properly, it must be applied after cutting the animal open along its dorsal midline but prior to spreading the larval body wall laterally (see dissection). The larva must be fixed with two minuten pins through the mouth hooks and the tail but without stretching the animal and without removing any inner organs yet. The preparation is bathed in the toxin solution for 10 minutes. After rinsing the preparation with HL3.1 saline, the body wall is spread laterally and dissected as described above (see dissection), and the specimen is mounted on the microscope stage ready for recording.

Synaptic transmission on muscle M6 in segment A2 and A3 was assessed in male animals carrying excisions of either *IS4A*, *I-IIA*, or *I-IIB* in cac on their only X-chromosome. Animals lacking exon *IS4B* are homozygous lethal. Thus, we used heterozygous females that expressed cac with *IS4B* excision on one X-chromosome over a cac^FLPStop^ construct which contains a tdTomato reporter followed by a stop codon within the open reading frame of all cac variants upon expression of a flipase transgene. Briefly, a reverse stop cassette followed by UAS-tdTomato flanked by canonical FRT sites of opposing orientation is inserted within a MiMIC residing in an intron of the Drosophila cac channel gene *cacophony*. Presence of canonical flipase turns the entire FlpStop-tdTomato cassette around and-re-inserts it in frame, which leads to pre-mature termination of cac transcription that is reported by tdTomato (Fisher et al. 2017). We expressed UAS-flipase under the control of the OK6 (Rapgap1)-GAL4 driver (Sanyal 2009), that drives expression of UAS-transgenes also in larval crawling motoneurons.

However, in our hands, flp events happen in a more or less stochastic manner. Flp events in motoneurons on M6 are less reliable as compared to Flp events on M12. As it is necessary that the FlpStop technique works in all motoneurons that innervate a target muscle to investigate the role of the IS4B exon for synaptic transmission, we used M12 (left and right) in A3 for this experiment rather than M6.

### Whole cell voltage clamp recordings

To investigate calcium currents that remain upon excision of IS4B, we used the same FlpStop approach as for synaptic transmission at the larval NMJ. UAS-flipase was expressed under the control of a DLM (dorsal longitudinal muscle) motoneuron Split-GAL4 fly strain that targets mainly the 5 wing depressor motoneurons (MN1-5) on each side of the body (10 neurons total, Hürkey et al. 2023). Expression of UAS-flipase under the control of this Split-GAL4 driver reliably led to expression of tdTomato in these motoneurons. In addition these flies are flight-less.

Adult MN patch clamp experiment were carried out as previously reported (Ryglewski et al. 2012). After dissection, the fly was positioned under a 40x water dipping lens (Zeiss, W Apochromat 40x/1.0 DIC VIS-IR ∞/0) on a fixed stage Zeiss AxioExaminer A1 upright fluorescence microscope. MN5 is situated on the dorsal surface of the ventral nerve cord in the mesothoracic neuromere. For whole cell voltage clamp recordings from MN5, protease was applied focally by alternating positive and negative pressure applied through a broken patch pipette to clean the membrane of the MN5 soma to facilitate seal formation. After cleaning, the preparation was perfused constantly with fresh saline (composition below). Voltage gated potassium channels were blocked by TEA, 4-AP and cesium in the recording solution (composition below). Tetrodotoxin (TTX, 10^-7^ M) to block voltage gated sodium channels was applied directly to the recording chamber while perfusion was halted for 3 minutes before approaching the cell with the patch pipette. Then perfusion resumed. Patch pipettes were filled with internal patch solution (composition below) and had resistances of ∼ 3.5 – 4 MΩ with this combination of internal and external solution. Recordings were carried out with an Axopatch 200B patch clamp amplifier (Molecular Devices). Data were digitized at 50 kHz (Digidata 1440, Molecular Devices) and filtered at 5 kHz through a lowpass Bessel filter. Data were recorded with pClamp 10.7.0.3 software package (Molecular Devices). After giga seal formation, whole cell configuration was achieved by a brief application of negative pressure to the patch pipette. We allowed 2 minutes for solution exchange with the cell interior and stabilization of the recording before adjusting whole cell capacitance and series resistance compensations and setting prediction at around 80% and correction at around 45%. Holding potential was –70 mV between voltage clamp protocols. Voltage gated calcium currents were elicited by voltage command steps to +20 mV in 10 mV increments from a holding potential of –90 mV. Leak was subtracted offline.

### Dissection and electrophysiology salines

Adult dissection saline [mM]: NaCl 128, KCl 2, CaCl_2_ 1.8, MgCl_2_ 4, HEPES 5, Sucrose ∼35.5. Osmolality was adjusted to 300 mOsM/kg with sucrose. pH was adjusted to 7.24 with 1N NaOH.

Larval dissection saline HL3.1 calcium free [mM]: NaCl 70, KCl 5, MgCl_2_ 4, NaHCO_3_ 10, trehalose 5, sucrose 115, HEPES 5. Osmolality was adjusted to 300 mOsM/kg with sucrose. pH was adjusted to 7.24 with 1N NaOH.

TEVC recording saline HL3.1 [mM]: NaCl 70, KCl 5, CaCl_2_ 0.5, MgCl_2_ 4, NaHCO_3_ 10, trehalose 5, sucrose 115, HEPES 5. Osmolality was adjusted to 300 mOsM/kg with sucrose. pH was adjusted to 7.24 with 1N NaOH.

Whole cell voltage clamp saline [mM]: NaCl 93.2, KCl 5, MgCl_2_ 4, CaCl_2_ 1.8, BaCl_2_ 1.8, Tetraethylammonium (TEA) Cl 30, 4-aminopyridine (4-AP) 2, HEPES 5, sucrose ∼35.5. Osmolality was adjusted to 300 mOsM/kg with sucrose. pH was adjusted to 7.24 with 1N NaOH.

Tetrodotoxin (10^-7^M) in normal saline was applied directly to the recording chamber.

Whole cell voltage clamp intracellular solution [mM]: CsCl 144, CaCl_2_ 0.5, EGTA 5, HEPES 10, TEA-Br 20, 4-AP 0.5, Mg-ATP 2. Osmolality was adjusted to 300 mOsm/kg with glucose. pH was adjusted to 7.24 with 1N CsOH.

### mEOS4b-tagged cac channel counting

The calculation of cac channel splice form numbers in individual AZs of the NMJ are based on the photoconversion of the fluorescent tag mEOS4b fused to the N-terminus of the cac channel (cac::mEOS4b-N; Ghelani et al. 2023). Third instar larvae were used for a body wall dissection. This preparation was suitable to image individual AZs within single boutons of an NMJ. We focused on muscle 6/7 on large type Ib boutons (Jia et al. 1993). Experiments were conducted at 25°C with HL3.1 solution (composition see above). We used a TIRF setup based on an inverted microscope (Nikon Eclipse Ti) equipped with a 60x 1.49 NA oil immersion objective (Nikon). Image series of up to 5000 frames were acquired at a frame rate of 20 Hz, using a sCMOS camera (Hamamatsu, Orca flash 4.0) controlled by the NIS-Element acquisition software (Nikon). Labelling of the NMJ with an anti-HRP antibody (diluted 1:1000) directly tagged with Alexa-488 for 5 min was used to have an initial landmark for NMJs within the preparation. During imaging of cac channels, we use a 1.5 magnification lens to reduce the effective pixel size to 71 x 71 nm. The used illumination protocol started with an initial continuous illumination with a 561 nm laser (80 % of initial laser power of 100 mW) for 250 frames to reduce the autofluorescent background. Afterward, we triggered the photoconversion of mEOS4b by switching on the photoconversion with a 405 nm UV-laser (5% of initial laser power of 100 mW) in addition to the 561 nm laser. The UV-excitation was sufficient to convert the majority of mEOS4b molecules into the red fluorescent state that was read out by the 561 nm laser. With the dual excitation of 405 nm and 561 nm, synapses were imaged until complete bleaching of the cac::mEOS4b-N population. The ability of the TIRF setup to adjust the laser beam orientation in the specimen was used to obtain an oblique illumination profile that limited the contribution of out of focus fluorescence. The number of channels was calculated as ratio of the maximal fluorescent intensity shortly after starting the photoconversion and the fluorescence of single mEOS4b molecules, which occurred as stochastic blinking events in the second half of the illumination sequence (see Fig. 4G, Ghelani et al. 2023). Two-dimensional x-y movements of the NMJ was corrected by using the NanoJ-Core drift correction from the NanoJ-Plugin for ImageJ/Fiji (Laine et al. 2019). After background subtraction, regions of 5×5 pixels were used to read out the fluorescence intensity profile of individual AZs. At least 30 AZs per ROI and larva were analyzed. For plotting the data and statistical calculations, we used Igor Pro 8 and GraphPad Prism 10.2.3.

### Immunohistochemistry

For cacophony immunohistochemistry, L3 larvae were dissected as described above and the VNC along with the motor nerves were removed. The specimen was then fixed for 7 minutes in ice cold (−30°C) 100% ethanol. For this, the saline was replaced by ice cold ethanol and rinsed a few times to assure cooling down of the specimen. Then the preparation was kept in the freezer at –30°C for 7minutes. Afterwards, the specimen was treated with PBS 0.1M with 0.3 % TritonX (PBS-Tx) at room temperature for 3×10 minutes, rocking.

cac^sfGFP^ (Gratz et al. 2019) localization (Figs. 2 and 3): Primary antibodies (α-brp nc-82, 1:400, DSHB; α-HRP rabbit, 1:500, Jackson Immunoresearch Cat# 323-005-021) were applied in PBS-Tx 0.3 % overnight at 4°C, rocking. Next day, primary antibody solution was removed, the preparation rinsed a few times with PBS-Tx 0.3 % and then washed 3×10 minutes with PBS-Tx 0.3 % at room temperature, rocking. 2 hour application of α-GFP nanobody (Fluotag X4 α-GFP Alexa 647, Nanotag Biotechnologies, Germany, Cat# N0304-AF647) and secondary antibodies donkey α-mouse Alexa 555 (Jackson Immunoresearch, Cat# 715-165-151) and donkey α-rabbit Alexa 488 (Jackson Immunoresearch, Cat# 711-545-152), all at a concentration of 1:500 at room temperature.

Assessment of ΔI-IIA and ΔI-IIB channel expression: Primary antibodies (α-brp nc-82, 1:400, DSHB; α-mEOS rabbit, 1:500, Badrilla, Cat# A010-mEOS2) were applied in PBS-Tx 0.3 % overnight at 4°C, rocking. Next day, primary antibody solution was removed, the preparation rinsed a few times with PBS-Tx 0.3 % and then washed 3×10 minutes with PBS-Tx 0.3 % at room temperature, rocking. 2 hour application of α-GFP nanobody (Fluotag X4 α-GFP Abberior Star Red, Nanotag Biotechnologies, Germany, Cat# N0304-ABRED) and secondary antibodies donkey α-mouse Alexa 555 (Jackson Immunoresearch, Cat# 715-165-151) and donkey α-rabbit Alexa 488 (Jackson Immunoresearch, Cat# 711-545-152), all at a concentration of 1:500 at room temperature.

For GluRIIA^GFP^ immunostaining, L3 larvae were fixed for 5 minutes in Bouin’s solution at room temperature. Then 3×10 minute washed were performed with 0.3% PBS-Tx at room temperature. This was followed by primary antibody incubation with α-brp (nc-82, 1:400, DSHB) overnight-After 3 more 10 minute washes with PBS-Tx 0.3%, specimen were incubated with α-GFP nanobody (Fluotag X4 α-GFP Abberior Star Red, Nanotag Biotechnologies, Germany, Cat# N0304-ABRED) and secondary donkey α-mouse Alexa 555 antibody (Jackson Immunoresearch, Cat# 715-165-151) for 3 hours at room temperature, covered to protect from light.

All immunolabel: Preparations were covered to prevent bleaching of fluorophores. Then, antibodies were removed, the preparations were rinsed a few times, which was followed by 3×10 minutes PBS (no Tx). Then an ascending ethanol series with 10 minues each with 50, 70, 90, 100 % ethanol was applied at room temperature, rocking. Preparations were then mounted in methylsalicylate, covered with a high precision cover slip that was sealed with clear nail polish. Preparations were kept in the dark until scanning with a Leica TSC SP8 upright confocal laser scanning microscope with a 40x oil NA 1.3 oil lens. Overviews were scanned without zoom at 1 µm z-step size. Magnifications were scanned with a 3.5 zoom with z-steps of 0.3 µm. All scans were done with a 3x line average.

### Assessment of fluorescence intensity

For fluorescence intensity measurement of sfGFP tagged cac channels in larval AZs, immunohistochemical labeling was carried out as for localization of cac channels. The only difference was that each round contained animals of all genotypes that were assessed. Animals were treated in the identical dish at the same time to ensure identical treatment. Confocal images were acquired with identical settings (lasers, detectors, scanning speed).

Fluorescence intensity was compared between genotypes by employing a custom Python script (available at doi://10.5281/zenodo.11299005) in which we used brp labeling (nc-82 antibody, DSHB) as a mask and asked for cac intensity within this mask. Intensity was then normalized to control. brp intensity was assessed identically.

GluRIIA fluorescence intensity was assessed identically, but as GluRIIA fields are larger than brp, we inflated the brp mask to encompass GluRIIA label. We then used the aforementioned python script to quantify intensity. For GluRIIA label we used flies that expressed a GluRIIA^GFP^ transgene that was previously shown to exhibit native GluRIIA expression levels (Rasse et al. 2005)

### Colocalization of cac and brp scaffold protein

To determine the Pearson’s correlation coefficient and the Manders co-occurrence coefficients of cac and brp stainings using the Costes’ threshold calculation (Manders et al. 1993; Costes et al. 2004) a custom Python script was utilized (available at doi://10.5281/zenodo.11299005).

In image stacks triple stained for cac, brp and HRP that had been deconvolved with the software Huygens, HRP staining served as a mask in which the desired coefficients were calculated for cac and brp.

### Super resolution microscopy (STED) and deconvolution

STED images were acquired at a Leica Stellaris 8 STED system (*Leica microsystems*) equipped with a pulsed white light laser (WLL) for excitation ranging from 440 to 790 nm and a 775 nm pulsed laser for depletion. Samples were imaged with a 93x glycerol objective (*Leica,* HC APO 93x/1.30 GLYC motCORR). For excitation of the respective channels the WLL was set to 640 nm for FluoTag-X4 anti-GFP Atto647N (NanoTag Biotechnologies, Cat# N0304-At647N) or 580 nm for Abberior Star 580 conjugated FluoTag-X2 anti-mouse antibody (NanoTag Biotechnologies, Cat# N2702-Ab580). Emission were detected in between 650-710 nm for Atto647N and in between 590-640 nm for Abberior Star 580. STED was attained by using the 775 nm laser for both channels. Additionally a 3^rd^ confocal channel was acquired (HRP, not shown, labeled with rabbit α-HRP antibody, Jackson Immunoresearch Cat# 323-005-021) by using 488 nm light for excitation of Alexa488 (Jackson Immunoresearch, Cat# 711-545-152) and an emission band of 500 – 530nm. Scanning properties were set to a format of 1024×1024 pixels, optical zoom factor to 5 (x/y=24.44 nm, z=191.69 nm) and scanning speed to 400 lines per second. Detectors were operated in in photon counting mode for both channels by 3 times line accumulation. For _gated_STED, detector time gates were set to 0.5-6 ns for both channels.

Deconvolution of STED stacks was done with Huygens Essential (*Scientific Volume Imaging, The Netherlands,* http://svi.nl). Within the *Deconvolution wizard*, images were subjected to background correction. *Signal-to-noise ratio* was set to 20*. The Optimized iteration mode of the CMLE* was applied by using 40 Iterations.

### Behavioral experiments

Wandering L3 larvae were selected off vial walls for crawling experiments and starved in empty vials for 5 minutes. Afterwards, they were placed in the middle of a plasticine square on an agarose gel (1% in H_2_O) using a brush. This square itself was placed on a tracking table with an acrylic glass plate. Usually, 10 larvae of the same genotype were recorded at a time. Occasionally, less than 10 larvae were available, in which case fewer larvae were recorded. Dishes were freshly poured each day before starting experiments. To prevent the larvae from escaping the boundary, an electrically charged wire was placed inside the square boundaries. Crawling pictures were obtained with a camera (Basler acA2040-90µm) from below the glass plate at a frame rate of 4 Hz for 5 minutes via pylon viewer (Version 6). Since few larvae were able to dig into the gel through small holes, only larvae that stayed inside the arena for 5 minutes were included into analysis. Tracking data of larval crawling were analyzed via the free software FIMtrack (University of Münster, Germany). Parameters that were read out included (but were not limited to) number of stops, average and maximum speed. The full list of data that were read out is available (doi://10.5281/zenodo.11299005). For explanation of read-out parameters, consult the FIMtrack manual (University of Münster website).

### Statistics and figure generation

Statistical analysis was performed with GraphPad Prism 10.2.1 (395). Data were tested for normal distribution using the Shapiro Wilk test. Normally distributed data were compared using either Student’s T-test (for two groups) or one-way ANOVA (for more than two groups) with subsequent Sidak’s Test post-hoc test for multiple comparisons or Dunnett’s test for each test group against control (planned comparisons). For non-normally distributed data either Mann-Whitney U-test (for two groups) or Kruskal Wallis Anova (with more than two groups) with subsequent Dunn’s post-hoc test for multiple comparisons or for test groups against control (planned comparisons) were used.

Differences were considered significant if p ≤ 0.05 *, p ≤ 0.01 **, p ≤ 0.001 ***, P ≤ 0.0001 ****.

Diagrams were generated with GraphPad Prism 10.2.1 (395) or Microsoft Excel 365. Figures with immunohistochemical images were generated with LasX software (Leica Microsystems, Wetzlar, Germany), with Fiji (www.Fiji.sc), or with Amira (version 6.5, Thermo Scientific). Images and diagrams were imported into CorelDraw Graphics Suite 2022 (Corel Corporation) either as Tiff images or as enhanced meta files (.emf) for figure production. Final figures were exported as .tif files.

All data and script is available at doi://10.5281/zenodo.11383963. Cacophony exon out flies will be sent to the Drosophila Bloomington Stock Center upon publication.

## Acknowledgments

We thank Kerstin Birod for performing Western Blots, Kate O’Connor-Giles for providing cacophony^mEOS4b^ flies, Tayfun Göncü for participating in gathering STED data, Susanne Hornig for confocal recordings of co-expressed cacophony exon out variants and Jan Werner for GluRIIA immunohistochemistry. This work was supported by a DFG research grant to S. Ryglewski (RY117/3-2) and to the Johannes Gutenberg Light Microscopy Core facility (Leica Stellaris STED, DFG INST 247/1004-1 FUGG).

## Figures and Legends

**supplementary Figure 1.**
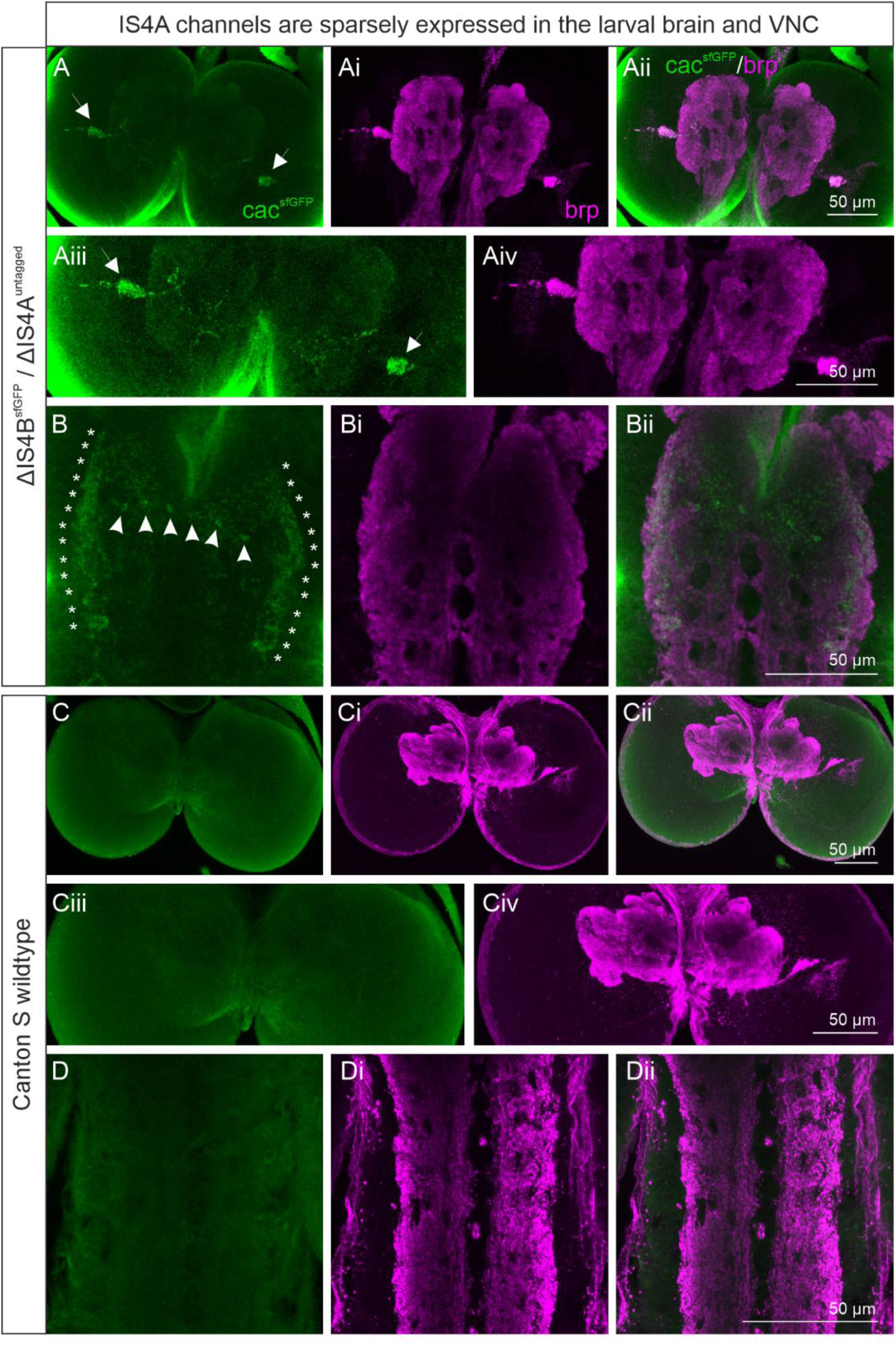
Cacophony^IS4A^ channels are sparsely expressed in the larval brain and ventral nerve cord. **(A, B)** Antibody labeling of GFP-tagged cacophony channels lacking exon IS4B (*ΔIS4B^sfGFP^*) reveals sparse cac label in the larval brain (green, **A-Aiv**, see arrows in **Aiii**) and the larval ventral nerve cord (green, **B-Bii**, see asterisks and arrow heads in B). Label was conducted in transheterozygous animals expressing cac^sfGFP^ ^IS4A^ over untagged cac^IS4B^ (*ΔIS4B^sfGFP^/ΔIS4A^no^ ^tag^*) to avoid expression of untagged IS4A channels. This likely increases labeling intensity. Cac label co-incides with AZ label as indicated by brp label (magenta, **Ai**, **Aiv**; **Bi**, **Bii**). **(C, D)** To exclude the possibility that cac label is misinterpreted due to overlap of with brp channel, antibody label was repeated in untagged Canton S wildtype animals **(C-CiV, D-Dii)**. Even strong brp label does not show in the green channel **(C, Ciii, D)**, thus indicating that cac^IS4A^ channels are indeed made and expressed. Images are maximum projections off F 13 single images, **(B)** 7 single images, **(C)** 10 single images, **(D)** 8 single images, z-size is 1 µm.

**supplementary Figure 2.**
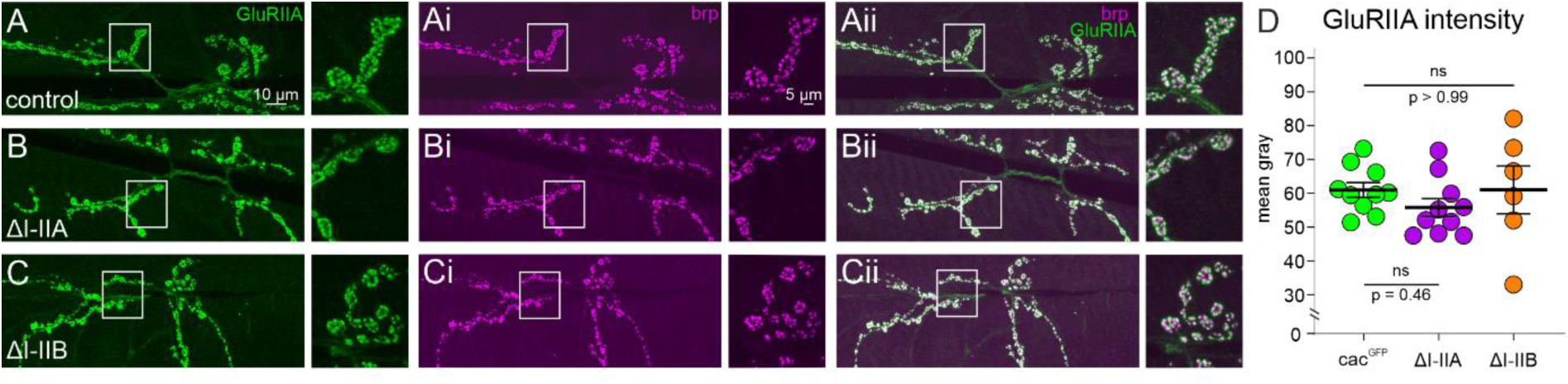
GluRIIA intensity across the NMJ is not altered upon excision of I-IIA or I-IIB. (**A-C**) GluRIIA is expressed in all NMJs independent of cacophony I-II exon excision (green; **A, Aii**: control, **B, Bii**: *ΔI-IIA*, **C, Cii**: *ΔI-IIB*, see overlap of GluRIIA and brp in **Aii-Cii**). The AZ scaffold protein brp (**Ai-Ci**) was used as mask to assess GluRIIA labeling intensity (**D**). Quantification of GluRIIA labeling intensity is shown as mean gray value and shows no difference between genotypes (**D**, cac^sfGFP^ control green, *ΔI-IIA^sfGFP^* purple, *ΔI-IIB^sfGFP^* orange, ANOVA with Dunn’s pairwise comparison).

